# Inhibition of nucleotide synthesis promotes replicative senescence of human mammary epithelial cells

**DOI:** 10.1101/423665

**Authors:** Alireza Delfarah, Sydney Parrish, Jason A. Junge, Jesse Yang, Frances Seo, Si Li, John Mac, Pin Wang, Scott E. Fraser, Nicholas A. Graham

**Affiliations:** Mork Family Department of Chemical Engineering and Materials Science, University of Southern California, University of Southern California, Los Angeles, 90089; Translational Imaging Center, Molecular and Computational Biology, University of Southern California, University of Southern California, Los Angeles, 90089; Norris Comprehensive Cancer Center, University of Southern California, University of Southern California, Los Angeles, 90089

**Keywords:** cellular senescence, epithelial cell, metabolomics, nucleoside/nucleotide biosynthesis, systems biology, growth arrest, cell stress, aging, ribonucleotide reductase regulatory subunit M2 (RRM2)

## Abstract

Cellular senescence is a mechanism by which cells permanently withdraw from the cell cycle in response to stresses including telomere shortening, DNA damage, or oncogenic signaling. Senescent cells contribute to both age-related degeneration and hyperplastic pathologies, including cancer. In culture, normal human epithelial cells enter senescence after a limited number of cell divisions, known as replicative senescence. Here, to investigate how metabolic pathways regulate replicative senescence, we used LC-MS–based metabolomics to analyze senescent primary human mammary epithelial cells (HMECs). We did not observe significant changes in glucose uptake or lactate secretion in senescent HMECs. However, analysis of intracellular metabolite pool sizes indicated that senescent cells exhibit depletion of metabolites from nucleotide synthesis pathways. Furthermore, stable isotope tracing with ^13^C-labeled glucose or glutamine revealed a dramatic blockage of flux of these two metabolites into nucleotide synthesis pathways in senescent HMECs. To test whether cellular immortalization would reverse these observations, we expressed telomerase in HMECs. In addition to preventing senescence, telomerase expression maintained metabolic flux from glucose into nucleotide synthesis pathways. Finally, we investigated whether inhibition of nucleotide synthesis in proliferating HMECs is sufficient to induce senescence. In proliferating HMECs, both pharmacological and genetic inhibition of ribonucleotide reductase regulatory subunit M2 (RRM2), a rate-limiting enzyme in dNTP synthesis, induced premature senescence with concomitantly decreased metabolic flux from glucose into nucleotide synthesis. Taken together, our results suggest that nucleotide synthesis inhibition plays a causative role in the establishment of replicative senescence in HMECs.

---

Cellular senescence is an irreversible growth arrest that occurs in response to a variety of cellular stresses including DNA damage, oncogenic signaling, and oxidative stress (1). Senescent cells accumulate in aging tissues and contribute to age-related decline in tissue function (2–4). In culture, replicative senescence occurs in normal cells after a limited number of population doublings, the so-called Hayflick limit (5). Following activation of oncogenes including KRAS and BRAF, senescence acts to prevent proliferation, thus preventing against cancer (6–9). In addition to its role in aging and tumor suppression, senescent cells can contribute to hyperplastic pathologies including cancer through secretion of numerous pro-inflammatory signals (ie, senescence-associated secretory phenotype, or SASP^1^) (10).

Although senescent cells are permanently arrested, they are highly metabolically active and demonstrate significant metabolic differences compared to proliferating, non-senescent cells (11). Studies of human fibroblasts in culture have shown that replicative senescence is accompanied by increased glycolysis (12–14). In oncogene-induced senescence, increased glucose consumption is shunted away from the pentose phosphate pathway, leading to decreased nucleotide synthesis (6, 15, 16). Alterations in mitochondrial function can also induce senescence through 5’ AMP-activated protein kinase (AMPK)- and p53-dependent pathways (17, 18). Importantly, it has also been demonstrated that metabolic genes including phosphoglycerate mutase, pyruvate dehydrogenase and malic enzymes can regulate entry into and escape from senescence (8, 19, 20). Therefore, identifying the mechanisms by which metabolism regulates senescence is essential to understanding the senescence program during aging and tumor suppression.

Primary human mammary epithelial cells (HMEC) have been shown to exhibit two mechanistically distinct senescence barriers to immortalization: stasis and agonescence (21). Stasis is a retinoblastoma-mediated growth arrest that occurs in the absence of DNA damage and is independent of p53 (22). Agonescence, or telomere dysfunction-associated senescence, is driven by critically shortened telomeres that trigger both a p53-dependent cell cycle arrest and a DNA damage response (23, 24). Because properties associated with senescence in mesenchymal cell types such as fibroblasts may not accurately reflect senescence in epithelial cells (22), the study of primary HMEC is required to understand how these senescence barriers are involved in normal HMEC cell biology, including aging and oncogenesis. How these senescence barriers are regulated by cellular metabolism in this primary cell type has not been previously investigated.

Here, we report that replicative senescence in primary human mammary epithelial cells (HMEC) is accompanied by a dramatic inhibition of nucleotide synthesis, including reduced flux of both glucose- and glutamine-derived carbon into nucleotide synthesis pathways. Expression of telomerase (hTERT) in HMEC both induced immortalization and maintained flux into nucleotide synthesis. In addition, treatment of proliferating HMEC with an inhibitor of ribonucleotide reductase regulatory subunit (RRM2), a key enzyme in dNTP biosynthesis, induced senescence and recapitulated the metabolomic state of replicative senescence. Taken together, our results indicate that nucleotide metabolism is a key regulator of replicative senescence in HMEC.

## RESULTS

### Establishment of a human mammary epithelial cell model of senescence

To study the metabolic alterations that accompany replicative senescence, we used normal diploid human mammary epithelial cells (HMEC). These cells have been previously shown to accurately represent the molecular changes that occur during replicative senescence *in vivo* (21). We observed linear growth for approximately 15 population doublings (PD) after which cell growth slowed until cells ceased proliferation at approximately 40 PD (Fig. 1A). HMEC at ~40 PD were viable and showed little to no cell death (Fig. 1B). However, these cells showed an enlarged, flattened and irregular morphology that is typical of senescent cells (25) (Fig. 1C). To confirm that HMEC at ~40 PD were senescent, we first measured senescence-associated beta-galactosidase (SA-β-gal) activity using C_12_FDG, a fluorogenic substrate for SA-β-gal activity in live cells (26) and observed increased SA-β-gal activity at PD 40 (Fig. 1D). Next, we confirmed that senescent HMEC exhibited a lack of DNA synthesis by measuring incorporation of the thymidine analog 5-ethynyl-2’-deoxyuridine (EdU) (Fig. 1E). Notably, Hoechst staining revealed that senescent HMEC did not exhibit senescence-associated heterochromatic foci (SAHF), a frequent but not obligatory marker of senescence (27). However, we observed that a significant fraction of senescent HMEC were multi-nuclear (Fig. 1F and Supporting Fig. 1), which has been observed in senescent human melanocytes systems (28) and can result from aberrant mitotic progression in oncogene-induced senescence (29). Next, we tested for expression of the senescence markers p16 and p21. Western blotting revealed that senescent HMEC exhibited increased expression of p21 but not p16 (Fig. 1G). Because SAHF formation is driven by p16 but not p21, this observation is consistent with our finding that senescent HMEC do not exhibit SAHF (27, 30). We also tested for expression of the senescence marker PAI-1 (plasminogen activator inhibitor-1) (31–33) by Western blot and observed increased expression in senescent HMEC (Fig. 1G). Finally, to investigate if senescent HMEC experienced DNA damage, we tested for expression of the DNA damage marker γ-H2AX by Western blotting. Interestingly, expression of γ-H2AX was slightly decreased as cells entered senescence (Fig. 1H) suggesting that senescent HMEC cells do not exhibit double-stranded DNA breaks (DSBs) (22). Taken together, these data demonstrate that HMEC enter senescence around 35-40 PD.

**Figure 1.**
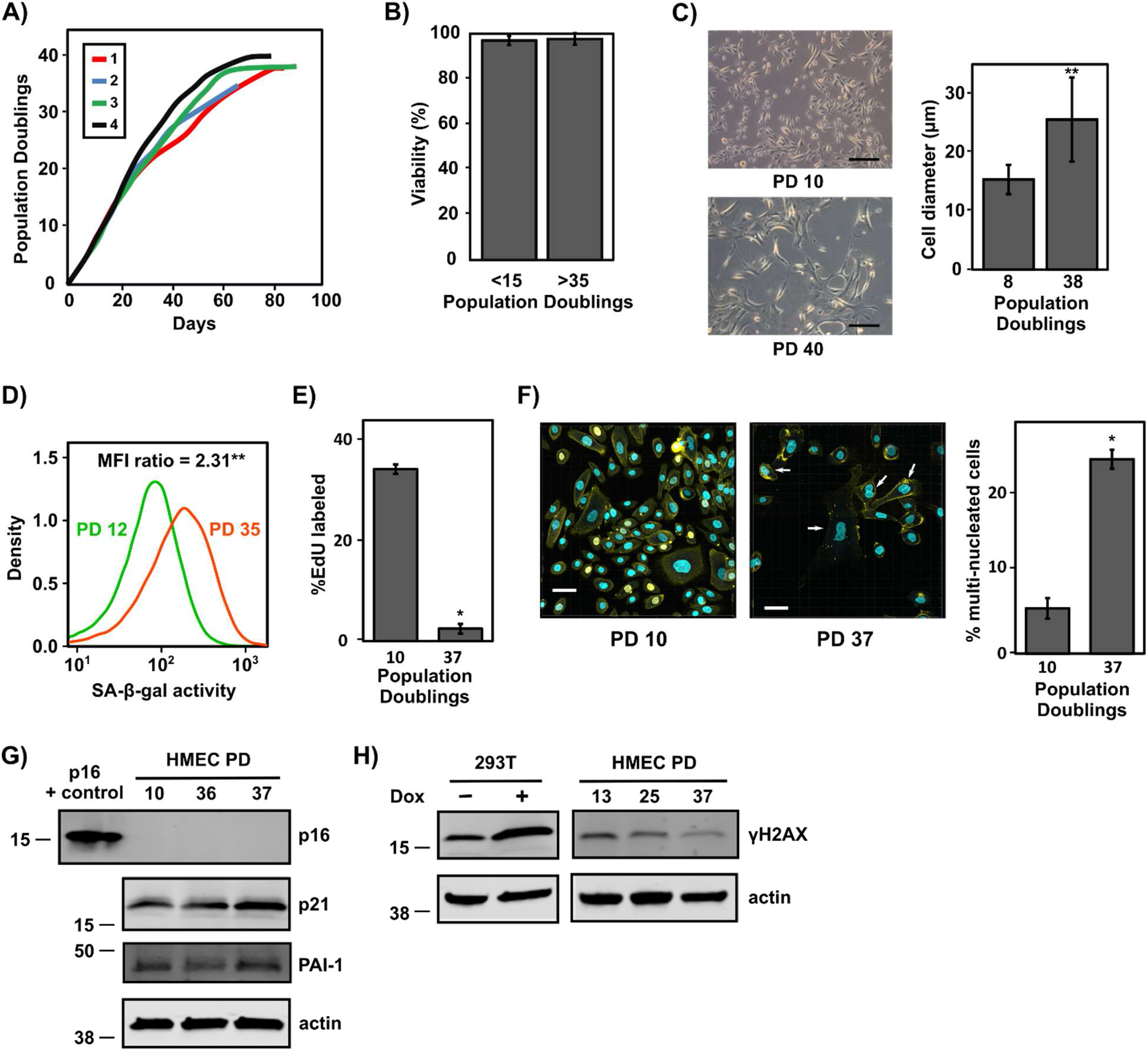
Establishment of an HMEC model of senescence. A) HMEC ceased proliferation at 35-40 population doublings. HMEC were cultured in M87A media with media replacement every 2 days (every day after 50% confluency) and passaged at 80-90% confluency (21). Four independent cultures are shown. B) HMEC viability was unaffected by senescence arrest. Viability of proliferating (<15 population doublings) and senescent (>35 population doublings). HMEC viability was measured by trypan blue staining. C) HMEC acquired enlarged, flattened and irregular morphology at PD 40. Phase contrast images and cell sizes represented at PD 10 and 40. Scale bar is 100 µm. ** denotes p-values less than 2×10^−16^ by Mann-Whitney U-test. D) HMEC showed increased activity of SA-β-gal at PD 35 measured by fluorescence signal of C_12_FDG. SA-β-gal measurements are shown at PD 10 and 35. SA-β-gal activity was calculated as (mean of samples labelled with C_12_FDG - mean of samples without C_12_FDG). ** denotes p-value less than 0.0001 by Student’s t-test. E) Measurement of DNA synthesis by EdU incorporation showed decreased DNA synthesis in senescent cells. ** denotes p-value less than 0.001 by Student’s t-test. F) Immunofluorescence imaging of proliferating and senescent HMEC using Hoechst and membrane dye. HMEC at PD 37 showed increased number of multi-nucleated cells (denoted by arrow). Scale bar is 50 µm. * denotes p-value less than 0.002 by Student’s t-test. G) Immunoblot for p16, p21, PAI-1, and actin with lysates from proliferating and senescent HMEC. Recombinant human p16 protein (40 ng) was used as a positive control in the p16 Western blot. H) Immunoblot for γ-H2AX and actin with lysates from proliferating and senescent HMEC. A lysate from 293T cell line treated with the DNA damaging agent doxorubicin (1 µM for 24 h) or control was used as a positive control. Actin was used as an equal loading control.

### Senescent HMEC do not exhibit a glycolytic shift

Having established a model system for replicative senescence, we analyzed proliferating and senescent HMEC using LC-MS-based metabolomics (34, 35). First, we measured the consumption and secretion rates of glucose and lactate, respectively (Fig. 2A, Supporting Table 1). Neither glucose consumption, lactate secretion, or the ratio of glucose consumption to lactate secretion was significantly altered, suggesting that senescent HMEC do not exhibit an overall glycolytic shift. Because senescent fibroblasts increase secretion of citrate (13), we next examined the secretion of TCA cycle metabolites. Senescent HMEC exhibited decreased secretion of most TCA cycle metabolites, though only aconitate and malate demonstrated significant reductions (Supporting Fig. 2A). Finally, we examined the consumption and secretion of amino acids in proliferating and senescent cells. Although consumption and secretion of many amino acids was reduced in senescent HMEC, only aspartate secretion was significantly reduced compared to proliferating HMEC (Supporting Fig. 2B). Notably, glutamine consumption, a significant carbon source for the TCA cycle, was not significantly reduced. Taken together, this data demonstrates that senescent HMEC are highly metabolically active, without significant changes in glycolytic ratio or in exchange of TCA cycle intermediates and amino acids with the extracellular medium.

**Figure 2:**
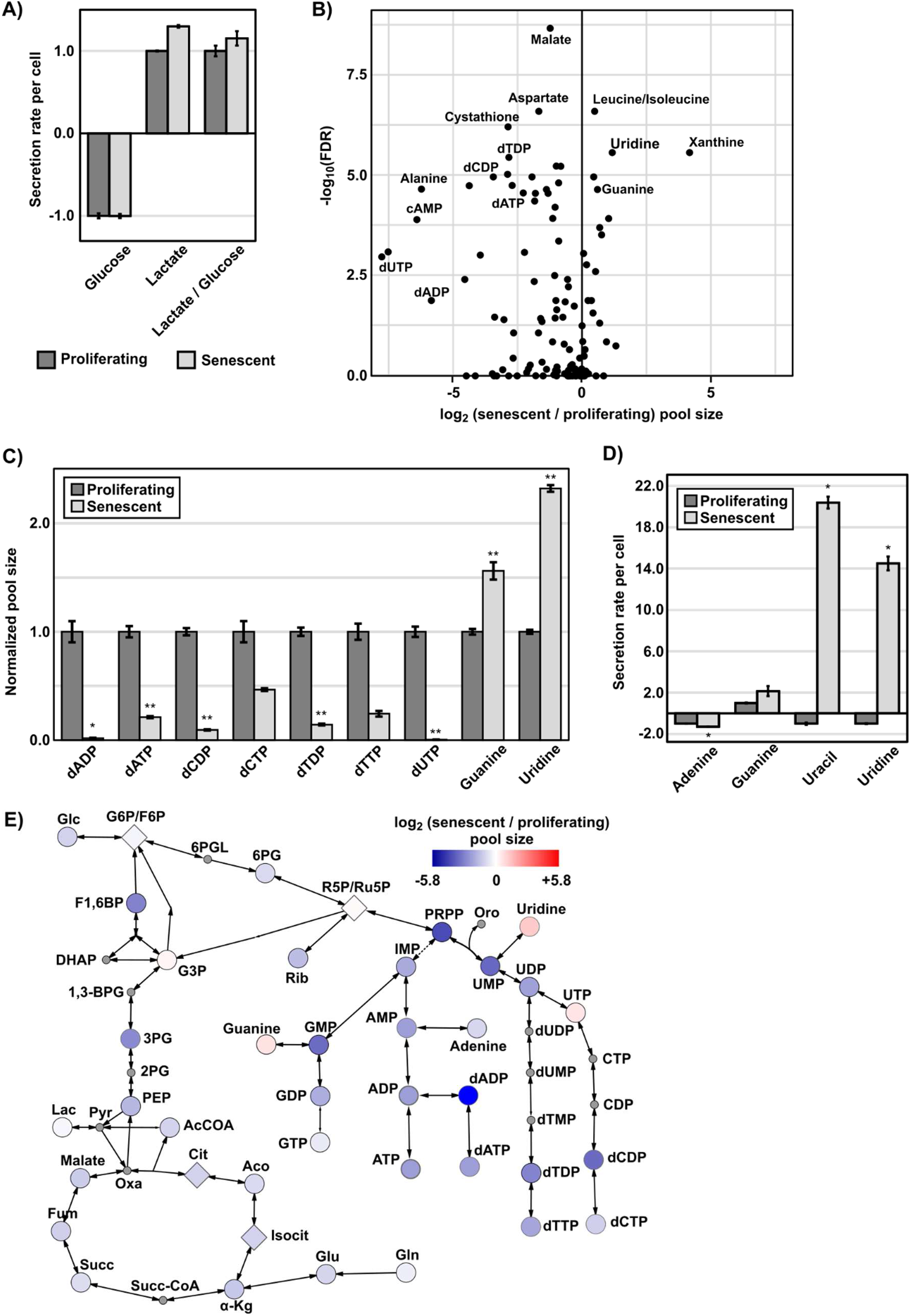
Replicative senescence in HMEC is accompanied by significant alterations in metabolite pool sizes. A) Glucose consumption, lactate secretion and ratio of glucose consumption to lactate secretion are unchanged in senescent HMEC. Metabolite extracts from blank and conditioned media were analyzed by LC-MS. Secretion or uptake is calculated as (conditioned media – blank media) normalized to integrated cell number. Secreted metabolites have positive values, and consumed metabolites have negative values. B) Volcano plot of intracellular pool sizes for all measured metabolites. Data represents average weighted log_2_ fold change (senescent/proliferating) and FDR-corrected Fisher’s combined p-value from five independent experiments. Depicted metabolites were measured in at least two independent experiments. C) Depletion of dNDPs and dNTPs and accumulation of nucleobases and nucleosides in senescent HMEC. Same data as in B for selected dNDPs, dNTPs, guanine and uridine. * and ** denote FDR-corrected Student’s t-test p-value less than 0.02 and 0.002, respectively. D) Extracellular secretion and consumption measured by LC-MS as in A for the nucleobases adenine, guanine, uracil and the nucleoside uridine. Secreted metabolites have positive values, and consumed metabolites have negative values. * denotes FDR-corrected Student’s t-test p-value less than 0.001. E) Metabolic pathway map depicting the log_2_ fold change (senescent/proliferating) intracellular pool sizes of metabolites in glycolysis, pentose phosphate pathway, nucleotide synthesis, and TCA cycle using the indicated color scale. Metabolites that were not measured are shown as small circles with grey color. Isomers that were not resolved with LC-MS are shown as diamonds.

### Replicative senescence in HMEC is accompanied by significant alterations in nucleotide and nucleoside pool sizes

Next, we quantified the intracellular metabolite pool sizes in both proliferating and senescent HMEC. In five independent experiments, we measured 111 metabolites in at least two independent experiments (Supporting Table 2). Of the measured metabolites, 11 were significantly increased and 31 were significantly decreased in senescent cells (false discovery rate-corrected p-value less than 0.01). Notably, we did not observe significant change in the AMP to ATP ratio of senescent HMEC (Supporting. Fig. 2C) in contrast to previous findings in senescent human fibroblasts (14, 36). Visualization of this data on a volcano plot revealed that several deoxyribonucleoside di- and tri-phosphates (dNDPs and dNTPs) including dCDP, dTDP, dADP, dATP, and dUTP were significantly downregulated in senescent cells (Fig. 2B,C). Conversely, intracellular levels of guanine and uridine were significantly upregulated. Increased intracellular levels of uridine were accompanied by increased secretion of uridine and the corresponding nucleobase uracil (Fig. 2D). Visualization of the log_2_ fold changes in intracellular pool size on a metabolic pathway map also revealed decreased metabolite pool sizes for most of glycolysis, the TCA cycle, pentose phosphate pathway and nucleotide synthesis metabolites (Fig. 2E). Metabolite set enrichment set analysis demonstrated that nucleoside mono/di/tri-phosphates were the most significantly downregulated metabolic pathway in senescent cells (Supporting Fig. 2D). Taken together, this data suggests that replicative senescence in HMEC induces dramatic changes in global metabolism with a particularly strong decrease in nucleotide metabolic pool sizes.

### Glucose-derived carbon fuels nucleotide synthesis in proliferating but not senescent HMEC

Because changes in metabolite pool size does not always accurately reflect changes in metabolic flux (37), we next analyzed metabolic flux in proliferating and senescent HMEC by stable isotope labeling. We cultured proliferating and senescent HMEC with [U-^13^C]-glucose followed by LC-MS metabolomics. To gain a global picture of the contribution of glucose-derived carbon to intermediary metabolites, we calculated glucose fractional contribution for each metabolite (38) (Supporting Table 3). Plotting the glucose fractional contribution on a volcano plot revealed that metabolites from the pyrimidine synthesis including UMP, UDP, UTP, dCDP, dCTP, dTDP, and dTTP showed significantly decreased incorporation of glucose-derived carbon (Fig. 3A and 3B). Notably, while phosphoribosyl pyrophosphate (PRPP), one of the precursors for pyrimidine synthesis, demonstrated only a small change in glucose-derived carbon incorporation, UMP and the downstream products of pyrimidine synthesis (eg, dCDP) exhibited almost complete loss of glucose-derived carbon incorporation (Fig. 3B and 3C). Visualization of the [U-^13^C]-glucose fractional incorporation on a metabolic pathway map confirmed that the pyrimidine synthesis pathway demonstrated the largest decrease in glucose fractional incorporation (Fig 3D). In contrast, metabolites from the TCA cycle showed little to no change in [U-^13^C]-glucose fractional incorporation or isotopomer distributions (Supporting Fig. 3).

**Figure 3:**
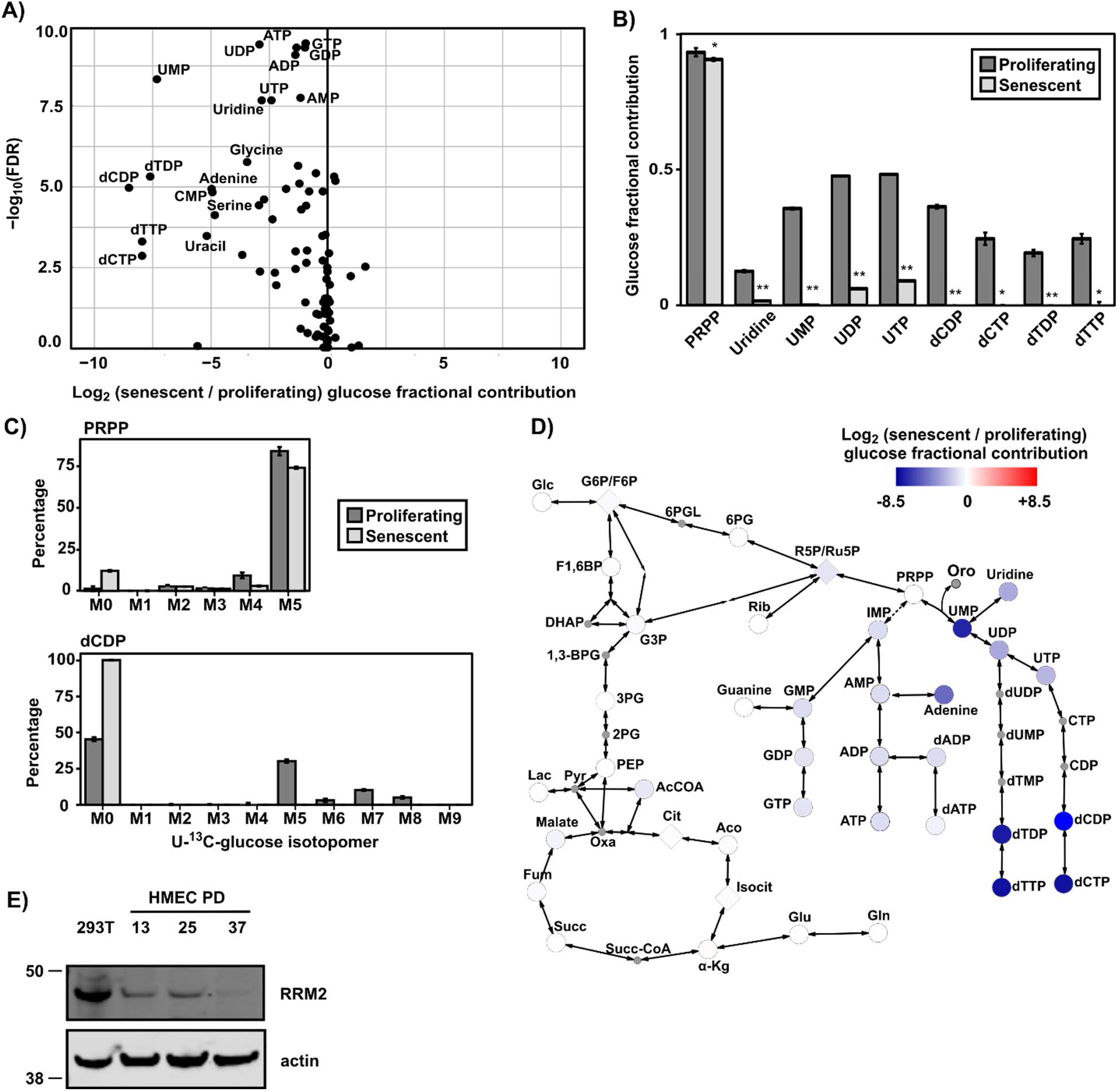
Glucose-derived carbon fuels nucleotide synthesis in young but not senescent HMEC. A) Purine and pyrimidine metabolites show significantly reduced [U-^13^C]-glucose carbon labeling in senescent HMEC. The volcano plot represents the average log_2_ fold change (senescent/proliferating) for fractional contribution (38) of [U-^13^C]-labeled glucose and FDR-corrected Fisher’s combined p-value from two independent experiments. B) Glucose flux into nucleotide synthesis pathways is globally reduced in senescent HMEC. Fractional contribution of [U-^13^C]-labeled glucose for selected pyrimidines. * and ** denote FDR-corrected Student’s t-test p-values less than 0.01 and 0.0001, respectively. C) [U-^13^C]-glucose isotopomer distributions for dCDP but not PRPP are significantly different in senescent HMEC. D) Metabolic pathway map depicting the average log_2_ fold change of fractional contribution from [U-^13^C]-glucose in senescent/proliferating HMEC on the indicated color scale for metabolites in glycolysis, pentose phosphate pathway, nucleotide synthesis, and the TCA cycle. Metabolites that were not measured are shown as small circles with grey color. Metabolites with isomers that were not resolved on LC-MS are shown as diamonds. E) Immunoblot for RRM2 and actin with lysates from proliferating and senescent HMEC. A protein lysate from 293T cell line was used as a positive control, and actin was used as an equal loading control. The RRM2 antibody was obtained from GeneTex.

We next used [1,2-^13^C]-glucose to investigate possible alterations in the ratio of glucose that enters the pentose phosphate pathway versus glycolysis (39). However, we observed no significant difference in the percentages of M1 and M2 lactate when comparing proliferating and senescent cells (Supporting Fig. 4A). There was, however, a significant decrease of ^13^C incorporation into UMP and downstream pyrimidines, consistent with our observation using [U-^13^C]-labeled glucose (Supporting Fig. 4B,C and Supporting Table 4). Next, we analyzed proliferating and senescent HMEC labeled with [U-^13^C]-glutamine to test for changes in TCA cycle flux. We observed decreased ratio of reductive to oxidative flux in the TCA cycle for senescent HMEC, however the fold change was small (Supporting Fig. 4D). Visualization of the [U-^13^C]-glutamine fractional contribution to metabolites revealed significant changes in contribution of glutamine-derived carbon into pyrimidine synthesis (Supporting Fig. 4E,F and Supporting Table 5). Taken together, this data suggests that senescent cells strongly downregulate the flux of carbon into pyrimidine synthesis while leaving glycolysis and the TCA cycle unaffected.

**Figure 4:**
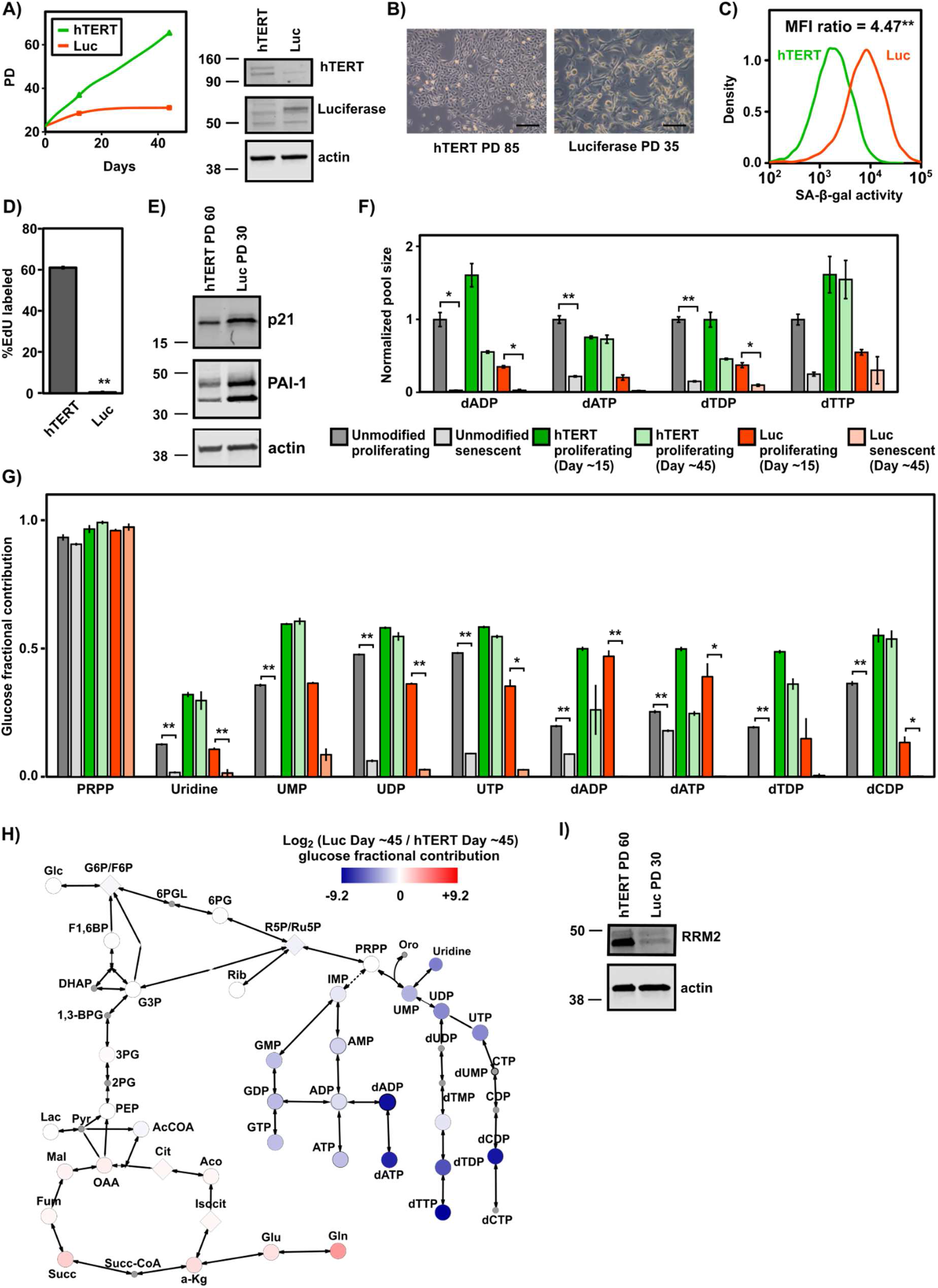
hTERT immortalization restores nucleotide pools and fluxes. A) Expression of hTERT in proliferating HMEC prevented occurrence of senescence growth arrest. A representative growth curve demonstrating that hTERT-expressing HMEC continued to grow logarithmically in culture while HMEC expressing luciferase ceased to proliferate at PD ~33. Filled triangles and squares indicate the PD at which cells were extracted for metabolomics. The x-axis represents days since completion of drug selection. hTERT and luciferase expression were confirmed by Western blotting. Actin was used as an equal loading control. B) Phase contrast images of luciferase- and hTERT-expressing HMEC at PD 35 and 85, respectively. Luciferase-expressing HMEC acquired enlarged, flattened and irregular morphology typical of senescent cells. hTERT-expressing HMEC maintained their epithelial cell morphology. Scale bar is 100 µm. C) Luciferase-expressing HMEC showed increased activity of SA-β-gal measured by fluorescence signal of C_12_FDG at SA-β-gal measurements are shown at Day ~60 following drug selection. SA-β-gal activity was calculated as (mean of samples labelled with C_12_FDG - mean of samples without C_12_FDG). ** denotes p-value less than 0.0001 by Student’s t-test. D) Measurement of DNA synthesis by EdU incorporation showed decreased DNA synthesis in senescent, luciferase-expressing cells at Day ~45 following drug selection. ** denotes p-value less than 0.00004 by Student’s t-test. E) Immunoblot for p21, PAI-1, and actin with lysates from luciferase- and hTERT-expressing HMEC at Day ~45 following drug selection. F) hTERT-expressing HMEC maintain purine and pyrimidine pools. Luciferase- and hTERT-expressing cells were profiled by LC-MS metabolomics at Days ~15 and ~45 following drug selection. Luciferase-expressing cells were senescent at Day ~45, but all other samples were still proliferating. Data from proliferating (PD 9) and senescent primary HMEC (PD 37) is shown for comparison. * and ** denote p-value less than 0.04 and 0.008, respectively. G) hTERT-expressing HMEC maintain glucose flux to purines and pyrimidines as measured by fractional contribution of [U-^13^C]-labeled glucose for selected purines and pyrimidines, as well as the nucleotide synthesis precursor PRPP. The number of days following drug selection for hTERT- and luciferase-expressing HMEC was the same as in F). Data from proliferating and senescent primary HMEC is shown for comparison. * and ** denote p-value less than 0.04 and 0.002, respectively. H) Metabolic pathway map depicting the average log_2_ fold change of [U-^13^C]-glucose fractional contribution for senescent, luciferase-(Day ~45) compared to proliferating, hTERT-expressing cells (Day ~45) on the indicated color scale for metabolites in glycolysis, pentose phosphate pathway, nucleotide synthesis, and the TCA cycle. Metabolites that were not measured are shown as small circles with grey color. Isomers that were not resolved with LC-MS are shown as diamonds. I) Immunoblot for RRM2 and actin with lysates from senescent, luciferase- and proliferating, hTERT-expressing HMEC.

We next queried published gene expression data to ask whether inhibition of nucleotide synthesis is broadly reflected at the transcriptional level in senescent cells. We analyzed i) microarray data from HMEC in replicative senescence (ie, stasis) (22); and ii) RNASeq data from IMR90 fibroblasts induced to senesce by ionizing radiation (2). Gene set expression analysis (GSEA) (40) across all KEGG metabolic pathways (n=78) (41) demonstrated a robust suppression of both pyrimidine metabolism (hsa00240) and purine (hsa00230) pathways in senescent cells (Supporting Fig. 5A,B and Supporting Table 6). In both data sets, expression of ribonucleotide-diphosphate reductase subunit M2 (RRM2), which catalyzes the biosynthesis of deoxyribonucleotides from ribonucleotides, was strongly decreased in senescent cells (not shown). Consistent with this data, Western blotting in our HMEC demonstrated downregulation of RRM2 in senescent cells (Fig. 3E and Supporting Fig. 5C). Taken together, this data suggests that senescent HMEC exhibit reduced nucleotide synthesis.

**Figure 5:**
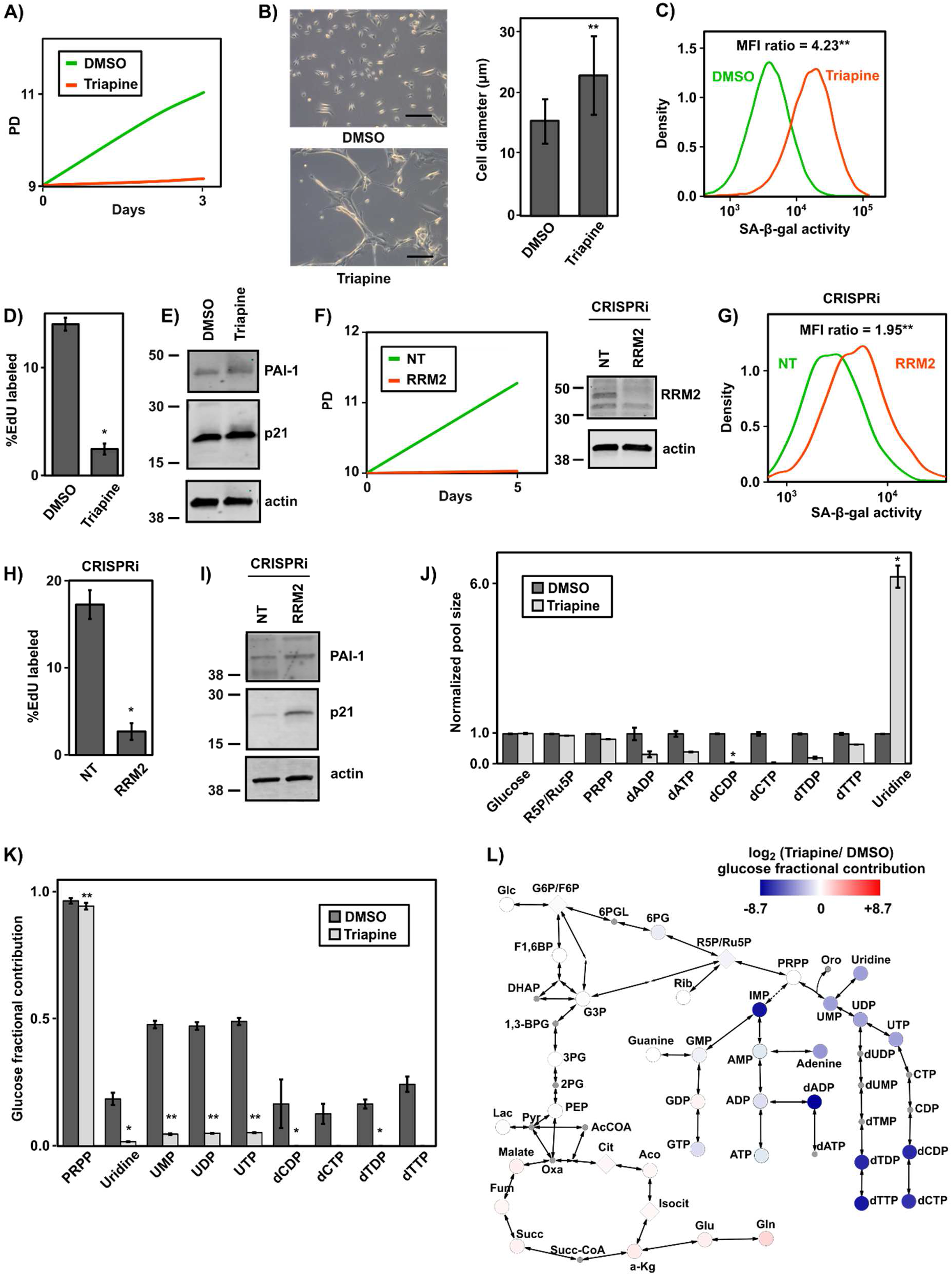
Inhibiting ribonucleotide reductase induces senescence. A) Treating proliferating HMEC with the RRM2 inhibitor triapine inhibits cell growth. Cells at PD ~8 were treated with 2 µM of triapine or DMSO for 72 h. B) Phase contrast images and cell sizes of HMEC treated with triapine or DMSO. HMEC acquire enlarged, flattened and irregular morphology typical of senescent cells within 3 days of treatment with triapine. Scale bar is 100 µm. ** denotes p-value less than 2×10^−16^ by Mann-Whitney U-test. C) Triapine-treated HMEC showed increased activity of SA-β-gal measured by fluorescence signal of C_12_FDG. SA-β-gal activity was calculated as (mean of samples labelled with C_12_FDG - mean of samples without C_12_FDG). ** denotes p-values less than 0.0001 by Student’s t-test. D) Measurement of DNA synthesis by EdU incorporation showed decreased DNA synthesis in triapine-treated cells. * denotes p-value less than 0.003 by Student’s t-test. E) Western blot of p21, PAI-1, and actin for triapine-treated cells F) Genetic knockdown of RRM2 expression inhibits HMEC growth. Cells at PD~10 were infected with sgRNA against RRM2 or non-targeting control. Western blot of RRM2 and actin for HMEC infected with sgRNA against RRM2 or non-targeting control. G) Knockdown of RRM2 expression induces SA-β-gal. SA-β-gal measured by fluorescence signal of C_12_FDG. SA-β-gal activity was calculated as (mean of samples labelled with C_12_FDG - mean of samples without C_12_FDG). ** denotes p-values less than 0.0001 by Student’s t-test. H) Measurement of DNA synthesis by EdU incorporation showed decreased DNA synthesis in RRM2 knockdown cells. * denotes p-value less than 0.02 by Student’s t-test. I) Western blot of p21, PAI-1, and actin for HMEC infected with sgRNA against RRM2 or non-targeting control. J) Triapine-treatment results in depletion of dNDPs and dNTPs and induces uridine accumulation. * denotes p-values less than 0.003 by Fisher’s combined t-test. K) Fractional contribution of [U-^13^C]-labeled glucose for selected pyrimidines. * and ** denote p-values less than 0.03 and 0.0001 by Fisher’s combined t-test, respectively. L) Metabolic pathway map depicting average log_2_ fold change of [U-^13^C]-glucose fractional incorporation for triapine/DMSO on the indicated color scale for metabolites in glycolysis, pentose phosphate pathway, nucleotide synthesis, and the TCA cycle. Metabolites that were not measured are shown as small circles with grey color. Isomers that were not resolved with LC-MS are shown as diamonds.

### HMEC immortalization with telomerase restores nucleotide pools and fluxes

Because transduction of HMEC with exogenous telomerase (hTERT) can immortalize pre-stasis HMEC (42), we next tested the effects of hTERT-mediated immortalization on the metabolic profiles of HMEC. We expressed either hTERT or a firefly luciferase control in proliferating HMEC (PD 13). hTERT-expressing HMEC continued to grow logarithmically in culture while HMEC expressing luciferase ceased to proliferate at PD ~33 (Fig. 4A). 60 days after transduction with luciferase and hTERT (PD 35 and 85 for luciferase and hTERT, respectively), we observed cuboidal cell morphology in hTERT-expressing cells, similar to proliferating, non-immortalized HMEC (Fig. 4B). In contrast, luciferase-expressing cells acquired a senescent morphology and exhibited increased SA-β-gal activity (Fig. 4B,C). Next, we confirmed that senescent luciferase-expressing HMEC exhibited a lack of DNA synthesis (Fig. 4D) and increased expression of the senescence markers p21 and PAI-1 (Fig. 4E). Taken together, this data demonstrates that hTERT expression efficiently immortalized HMEC.

We then profiled luciferase- and hTERT-expressing HMEC using LC-MS metabolomics. Consistent with unmodified HMEC, media footprint analysis revealed that glucose consumption and lactate secretion rates were not significantly different in proliferating, hTERT-expressing and senescent, luciferase-expressing cells (Supporting Fig. 6 and Supporting Table 7). Secretion of guanine and uridine was increased in senescent, luciferase-expressing HMEC. We next examined intracellular metabolite pool sizes in luciferase- and hTERT-expressing cells at Day ~15 and ~60 following viral transduction and drug selection. At Day ~15, both luciferase- and hTERT-expressing cells were proliferating, whereas at Day ~60, luciferase- but not hTERT-expressing cells were senescent. Metabolomic profiling demonstrated that hTERT-expressing cells experienced no significant changes in dNDP and dNTP pool sizes between Days ~15 and ~60 (Fig. 4F and Supporting Table 8). In contrast, luciferase-expressing cells exhibited significantly decreased dNDP and dNTP metabolite pool sizes, closely mirroring observations in untransduced proliferating and senescent HMEC.

**Figure 6:**
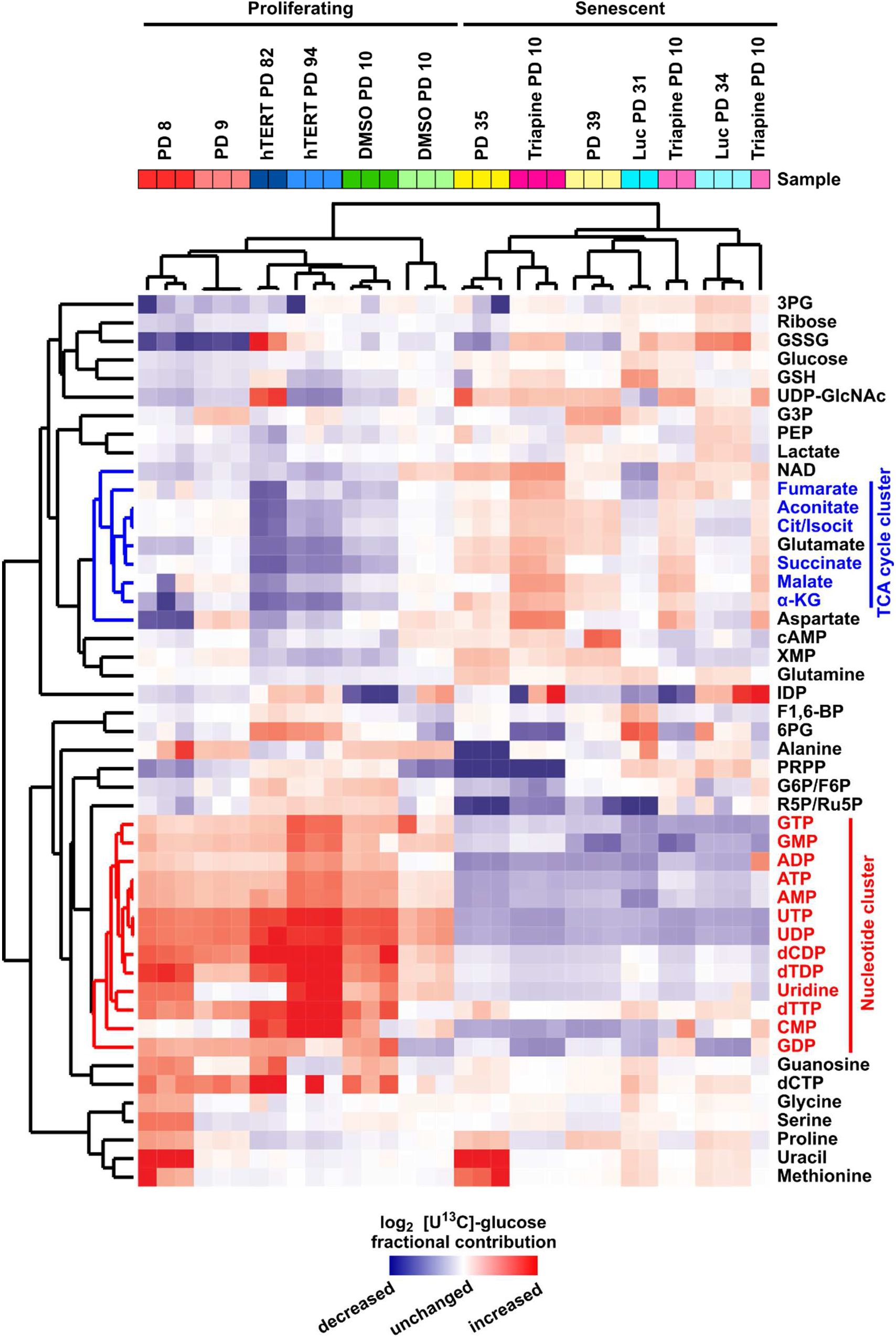
Integrative analysis of glucose contribution to metabolites in proliferating and senescent HMEC. Heatmap comparing fractional contribution of glucose to commonly measured metabolites. Columns represent individual samples. Red indicates upregulation and blue indicates downregulation. Nucleotides cluster together and are upregulated in proliferating samples compared to senescent samples.

Next, we used [U-^13^C]-labeled glucose to assess the fractional contribution of glucose to the metabolism of HMEC expressing hTERT and luciferase. This analysis revealed that [U-^13^C]-glucose fractional incorporation in hTERT-expressing cells was unchanged between Days ~15 and ~60 for pyrimidines and purines including UMP, UDP, UTP, dADP, dATP, dTDP, and dCDP (Fig. 4G, Supporting Fig. 7, and Supporting Table 9). In contrast, luciferase-expressing HMEC exhibited significantly reduced fractional incorporation of [U-^13^C]-glucose into these pyrimidines and purines, similar to observations in untransduced proliferating and senescent HMEC.

Visualization of the glucose fractional contribution on a metabolic pathway map confirmed a global inhibition of carbon flux to nucleotides downstream of PRPP (Fig. 4H). In addition, metabolite set enrichment set analysis confirmed that nucleoside mono/di/tri-phosphates exhibited significantly decreased [U-^13^C]-glucose-derived label in senescent, luciferase-expressing HMEC (Supporting Fig. 8). Notably, hTERT also induced moderate downregulation of glucose-derived carbon to TCA cycle metabolites, a difference that was not observed comparing unmodified proliferating and senescent HMEC. Finally, we measured protein expression of RRM2 using western blot and observed significant decrease in high PD senescent luciferase-expressing HMEC (Fig. 4I). Taken together, these results demonstrate that hTERT-immortalization maintains nucleotides pools and fluxes in HMEC.

### Inhibiting ribonucleotide reductase induces senescence

Based on the metabolomic profile of senescent and proliferating HMEC, we hypothesized that reduced nucleotide synthesis plays a causative role in induction of senescence. To test this hypothesis, we first treated proliferating HMEC with triapine, an inhibitor of ribonucleotide reductase regulatory subunit (RRM2) (43). Indeed, treatment of proliferating HMEC (PD 10) with triapine effectively blocked cell proliferation (Fig 5A). Following triapine treatment, cells acquired a typical senescent morphology and were significantly increased in size (Fig. 5B). Additionally, triapine-treated HMEC exhibited increased SA-β-gal activity (Fig. 5C), reduced DNA synthesis (Fig. 5D), and increased expression of p21 and PAI-1 (Fig. 5E and Supporting Fig. 9A). To confirm the role of RRM2 in senescence, we next used CRISPRi-mediated knockdown of RRM2 expression (CRISPR-dCas9-KRAB) (44). Expression of a sgRNA against RRM2 but not a non-targeting sgRNA control efficiently knocked down RRM2 expression and inhibited cell growth in proliferating HMEC (Fig. 5F). We then confirmed that RRM2 knockdown increased SA-β-gal activity (Fig. 5G), reduced DNA synthesis (Fig. 5H), and increased expression of p21 and PAI-1 (Fig. 5I and Supporting Fig. 9B). Taken together, these data demonstrate that inhibition of RRM2 is sufficient to induce senescence in primary HMEC.

Next, we tested whether inhibition of RRM2 could induce senescence in the immortalized but non-transformed mammary epithelial cell line MCF10A. Indeed, triapine treatment slowed cell growth, induced SA-β-gal activity, reduced DNA synthesis, and increased expression of p21 and PAI-1 (Supporting Fig. 9C-G). Together with data from primary HMEC, these data demonstrate that inhibition of RRM2 activity can induce senescence in immortalized mammary epithelial cells.

To investigate metabolomic alterations upon triapine-induced senescence, we again performed untargeted LC-MS metabolomics. HMEC treated with triapine exhibited depleted dNDP and dNTP pool sizes and increased intracellular levels of the nucleoside uridine (Fig. 5J). In contrast, we observed no significant change in the intracellular pool sizes of glucose, ribose-5-phosphate and PRPP (Fig 5J and Supporting Table 10). Tracing [U-^13^C]-labeled glucose in triapine-treated HMEC highlighted the downregulation of glucose flux into nucleotide biosynthesis (Fig. 5K). Examination of the [U-^13^C]-labeled glucose incorporation levels in triapine-treated cells showed that fractional contribution of glucose to PRPP was not significantly changed, similar to replicative senescence (Fig. 5K,L and Supporting Table 11). However, there was a significant inhibition of glucose flux into pyrimidines in triapine-induced senescent HMEC (Fig. 5K,L). In contrast, glucose flux into TCA cycle did not significantly change upon triapine treatment (Fig. 5L). Metabolite set enrichment set analysis demonstrated that nucleoside and deoxynucleoside mono/di/tri-phosphates were the most significantly decreased metabolic pathways in triapine-induced senescent cells (Supporting Fig. 10). Taken together, these results indicate that inhibition of nucleotide synthesis is sufficient to trigger senescence in non-immortalized HMEC.

### Integrative analysis of glucose contribution to metabolites in proliferating and senescent HMEC

To identify the most consistent alterations in senescent cells across our metabolomic experiments, we conducted unsupervised hierarchical clustering of the glucose fractional contribution to each metabolite across experiments. Proliferating HMEC cultures (low PD, hTERT expression, and DMSO treatment) and senescent HMEC cultures (high PD, luciferase expression, and triapine treatment) were separated into two well defined clusters (Fig. 6). In addition, nucleotides were tightly clustered together to reveal decreased levels of glucose-derived carbon incorporation in senescent cells (red cluster). This heatmap also highlighted the moderate downregulation of glucose contribution to TCA cycle metabolites in hTERT-expressing HMEC (blue cluster). Taken together, this confirms that inhibited nucleotide synthesis is a hallmark of senescent HMEC.

## DISCUSSION

Cellular senescence is a state of irreversible cell cycle arrest that contributes to degenerative and hyperplastic phenotypes in aging and cancer. Although senescent cells are withdrawn from the cell cycle, they remain highly metabolically active (11). Here, we show that inhibition of nucleotide synthesis regulates replicative senescence in primary HMEC. The inhibition of nucleotide synthesis in senescent HMEC was reflected both in reduced pool sizes (Fig. 2E) and in reduced metabolic flux into nucleotide synthesis (Fig. 3D and Supporting Fig. 4). In contrast, fluxes from glucose and glutamine into glycolysis, the pentose phosphate pathway, and the TCA cycle were not significantly changed. Importantly, treatment of proliferating HMEC with an inhibitor of RRM2, a key enzyme in dNTP synthesis, demonstrated that inhibition of nucleotide synthesis was sufficient to induce senescence and recapitulate the metabolomic signature of HMEC replicative senescence. Taken together, these findings demonstrate the crucial role of nucleotide synthesis in cellular senescence of a primary epithelial cell type, a finding with implications for both aging and tumor development.

The study of metabolism during replicative senescence has relied heavily on fibroblasts, a mesenchymal cell type. These studies have demonstrated that human fibroblasts gradually exhibit a more glycolytic metabolism as they become senescent (12–14). Similarly, senescence induced by oncogene activation also leads to increased glycolysis (8, 16). Here, using a primary epithelial cell type (HMEC), we did not observe significant alterations in either glucose uptake or lactate secretion in senescent HMEC (Fig. 2A). Additionally, the fractional contribution of glucose and glutamine to glycolysis and the TCA cycle was unchanged in senescent HMEC. In contrast to senescent fibroblasts (14), HMEC senescence was also not accompanied by an energy crisis (Supporting Fig. 2C). Taken together, these results suggest that epithelial and fibroblast cells exhibit distinct metabolic alterations during senescence. These metabolic differences may reflect the fact that senescent epithelial and fibroblast cells exhibit distinct transcriptional profiles (22).

Although replicative senescence in our HMEC system does not involve oncogene activation, the inhibition of nucleotide synthesis we observed resembles the metabolic alterations found in oncogene-induced senescence. Notably, in fibroblasts, oncogenic RAS induces senescence by suppressing nucleotide metabolism, leading to the depletion of dNTP pools (6, 9). This depletion of dNTP pools is mediated by RAS-induced repression of RRM2 mRNA and protein levels. Similarly, senescent HMEC showed a significant downregulation of RRM2 protein expression (Fig. 3E and Supporting Fig. 5C). Notably, in HMEC, chemical inhibition of RRM2 was sufficient to induce senescence and recapitulated the metabolomic signature of senescent HMEC (Figs. 5 and 6). Together, our findings extend the relevance of nucleotide metabolism in senescence to a non-transformed, primary cell type in the absence of oncogene activation.

In the current study, we observed depletion of dNTPs in the absence of γ-H2AX induction, suggesting that senescent HMEC do not experience DNA double strand breaks during the time course of our experiments (Fig. 1E). However, because the maintenance of sufficient and balanced dNTP pools is essential for maintaining genomic integrity (45, 46), the inability to synthesize dNTPs may eventually lead to replication stress, genomic instability, and mutagenesis (47, 48). Indeed, genomic instability in the early stages of tumorigenesis induced by cyclin E oncogenes is coupled to insufficient nucleotide availability (49). In addition, we observed significantly increased number of multi-nucleated cells in the senescent HMEC culture which has previously been linked to genomic instability (50), telomere dysfunction (51), polyploidy (52) and tumor-initiation (28). Additionally, the extent of senescence experienced by mouse embryonic fibroblasts during immortalization correlates with the amount of genomic instability in immortalized cell lines (34). Interestingly, in RAS-expressing fibroblasts, senescence-associated nucleotide deficiencies can be bypassed by inactivation of the serine/threonine kinase ATM, which is activated by DNA damage (53). However, because senescent HMEC in this study are not experiencing DNA double strand breaks, they may more closely resemble hypoxic cells which experience replication stress but not DNA damage (54). Similar to hypoxic cells, senescent cells upregulate RRM2B (55), a subunit of the ribonucleotide reductase complex which may preserve the capacity for sufficient nucleotide synthesis during hypoxia to avoid DNA damage.

In senescent HMEC, we observed that the flux of glucose-derived carbon was more severely downregulated in pyrimidine synthesis than in purine synthesis (Figs. 3D, 4H, and 5L). This difference between pyrimidine and purine synthesis was not reflected in metabolite pool sizes, highlighting the power of stable isotope tracing to reveal metabolic changes not apparent in metabolite pool sizes (37, 56). Pyrimidine synthesis can be specifically controlled through mTOR/S6K1-mediated phosphorylation of CAD (carbamoyl-phosphate synthetase 2, aspartate transcarbamylase, and dihydroorotase), the enzyme that catalyzes the first three steps of *de novo* pyrimidine synthesis (57). Additionally, loss of urea cycle enzymes including carbamoyl phosphate synthetase-1 (CPS1) can lead to pyrimidine but not purine depletion (58). Although it is not yet clear why pyrimidines are more affected than purines in senescent HMEC, this observation remains a subject of active investigation.

Taken together, our results indicate that suppression of nucleotide synthesis is a critical metabolomic alteration regulating senescence in primary HMEC. A more detailed understanding of altered metabolism as both a cause and a consequence of senescence will have important implications. In both aging (2) and cancer (59), therapeutic targeting of senescence has shown great promise. Similarly, therapeutics targeting the emerging metabolic differences between senescent and non-senescent cells may prove useful tools for improving human health.

## EXPERIMENTAL PROCEDURES

### Cell culture

HMEC cells were purchased from Thermo Scientific and cultured in M87A medium (50% MM4 medium and 50% MCDB170 supplemented with 5 ng/ml EGF, 300 ng/ml hydrocortisone, 7.5 ug/ml insulin, 35 ug/ml BPE, 2.5 ug/ml transferrin, 5 µM isoproterenol, 50 µM ethanolamine, 50 µM o-phosphoethanolamine, 0.25 % FBS, 5 nM triiodothyronine, 0.5 nM estradiol, 0.5 ng/ml cholera toxin, 0.1 nM oxytocin, 1% anti-anti, no AlbuMax) in atmospheric oxygen. MCF10A cells were obtained from American Tissue Culture collection (ATCC) and cultured in DMEM/F-12 medium supplemented with 20 ng/ml EGF, 500 ng/ml hydrocortisone, 100 µg/ml insulin, 100 ng/ml cholera toxin, 0.1 nM oxytocin, 5% horse serum and 1X Pen/Strep in atmospheric oxygen. Glucose and glutamine-free DMEM was purchased from Corning (90-113-PB), Ham’s F12 was purchased from US Biological (N8542-12), and MCD170 medium was purchased from Caisson Labs (MBL04). Glucose and glutamine were added to the media at the appropriate concentration for each media type. Cells were lifted with TrypLE at 80-90% confluency and seeded at a density of 2.3× 10^3^/cm^2^. Cell viability and diameter was measured with trypan blue assay using TC20 automated cell counter (Bio-Rad).

### Genetic modification

Proliferating HMEC were infected at PD 14 with pLenti-PGK-hygro (Addgene 19066) encoding either hTERT or firefly luciferase. Following infection, cells were selected with 5 µg/ml hygromycin for 7 days. Following selection, cells were maintained in culture with 2 µg/ml hygromycin. To knockdown expression of RRM2, proliferating HMEC were infected at PD 10 with lentiviral vector expressing dCas9-KRAB and sgRNA against RRM2 (forward 5’-CACCGACACGGAGGGAGAGCATAG-3’) or non-targeting control (forward 5’-CACCGGTATTACTGATATTGGTGGG-3’). The lentiviral vector used for CRISPRi, pLV hU6-sgRNA hUbC-dCas9-KRAB-T2a-Puro, was a gift from Charles Gersbach (Addgene plasmid # 71236)

### Senescence-associated β-galactosidase measurements

HMEC were incubated with 100 nM bafilomycin A1 (Sigma-Aldrich) for 1 h to raise the lysosomal pH to 6.0 (26), followed by incubation with 33 µM C_12_FDG (Setareh Biotech) for 1 h, lifted with TrypLE, resuspended in ice-cold PBS, and then analyzed on a Miltenyi Biotec MACSQuant flow cytometer to measure fluorescence (26). Data was processed and analyzed with FlowJo 7.6.1 software and the mean fluorescent signal for each sample was exported. Values for SA-β-gal activity were calculated as (mean of samples labelled with C_12_FDG -mean of samples without C_12_FDG). Data was normalized to the SA-β-gal activity of non-senescent samples.

### EdU incorporation

EdU staining was performed using Click-iT Plus EdU Alexa Fluor 594 Imaging kit (Thermo Fisher, C10639). Briefly, cells were seeded in glass bottom dishes (MatTek Part No: P35G-1.5-10-C) with appropriate density and then incubated with 10 µM EdU for 6 h at 37 °C. If membrane staining desired, cells were incubated with 1X Membrite Fix 640/660 (Biotium 30097-T) for 5 mins at 37 °C. Next, cells were fixed with 3.7% formaldehyde followed by 0.5% Triton X-100 permeabilization at room temperature. Cells were incubated with Click-iT Plus reaction cocktail for 30 min at room temperature protected from light. Finally, for nuclear staining, cells were incubated with 1X Hoechst 33342 for 30 mins at room temperature protected from light and stored in PBS until imaging.

### Confocal microscopy and cell counting

All microscopy was performed on Zeiss 780 or 880 confocal systems. The excitation wavelengths for the stained cells were: 405 nm for Hoechst, 594 nm for EdU, and 633 nm for MemBrite membrane dye (Biotium). All high magnification images (40x/1.1NA and 63x/1.4NA, Zeiss) were sampled at Nyquist with 1 Airy Unit pinhole diameters and all low magnification images (10x/0.45NA, Zeiss) were collected with pinhole diameters such that 40 µm optical sections were obtained for cell counting. Tiled images were stitched together and analyzed using Imaris (Bitplane) image processing and analysis tools. Images were interrogated for nuclei and presence of EdU through the Spots module in Imaris.

### Mass spectrometry-based metabolomics analysis

HMEC were plated onto 6-well plates at density of 1-3 × 10^4^ cells/cm^2^ depending on the experiment. For flux analysis, after 24 h media was replaced by [U-^13^C]-labeled glucose, [1,2-^13^C]-labeled glucose, or [U-^13^C]-labeled glutamine (Cambridge Isotope Laboratories). Metabolite extraction was performed 24 h after adding labeled media. For extraction of intracellular metabolites, cells were washed on ice with 1 ml ice-cold 150 mM ammonium acetate (NH_4_AcO, pH 7.3). 1 ml of −80 °C cold 80% MeOH was added to the wells, samples were incubated at −80 °C for 20 mins, then cells were scraped off and supernatants were transferred into microfuge tubes. Samples were pelleted at 4°C for 5 min at 15k rpm. The supernatants were transferred into LoBind Eppendorf microfuge tube, the cell pellets were re-extracted with 200 µl ice-cold 80% MeOH, spun down and the supernatants were combined. Metabolites were dried at room temperature under vacuum and re-suspended in water for LC-MS run. For extraction of extracellular metabolites, 20 µl of cell-free blank and conditioned media samples were collected from wells. Metabolites were extracted by adding 500 µl −80 °C cold 80% MeOH, dried at room temperature under vacuum and re-suspended in water for LC-MS analysis.

Samples were randomized and analyzed on a Q Exactive Plus hybrid quadrupole-Orbitrap mass spectrometer coupled to an UltiMate 3000 UHPLC system (Thermo Scientific). The mass spectrometer was run in polarity switching mode (+3.00 kV/-2.25 kV) with an m/z window ranging from 65 to 975. Mobile phase A was 5 mM NH_4_AcO, pH 9.9, and mobile phase B was acetonitrile. Metabolites were separated on a Luna 3 µm NH_2_ 100 Å (150 × 2.0 mm) column (Phenomenex). The flowrate was 300 µl/min, and the gradient was from 15% A to 95% A in 18 min, followed by an isocratic step for 9 min and re-equilibration for 7 min. All samples were run in biological triplicate.

Metabolites were detected and quantified as area under the curve based on retention time and accurate mass (≤ 5 ppm) using the TraceFinder 3.3 (Thermo Scientific) software. Raw data was corrected for naturally occurring ^13^C abundance (60). Extracellular data was normalized to integrated cell number, which was calculated based on cell counts at the start and end of the time course and an exponential growth equation. Intracellular data was normalized to the cell number and cell volume at the time of extraction. Pathway maps were made with Cytoscape software (61).

### Gene set and metabolite Set enrichment Analysis

For gene set expression analysis of RNA data, microarray data from HMEC (22) was compared across pairwise comparisons for stasis and pre-stasis cell cultures. Genes were then ranked by their average rank from individual experiments. RNAseq data from IMR90 cells (2) was ranked based on signal to noise ratio of senescent/non-senescent cells. GSEA was run with the unweighted statistic using the GSEA java applet (40). For metabolite set enrichment analysis of metabolomic data, HMEC intracellular pool sizes or ^13^C-glucose fractional contribution data was ranked based on log_2_ fold change of senescent/proliferating, luciferase/hTERT or triapine/DMSO. Enrichment analysis was run with unweighted statistic using the GSEA java applet.

### Statistical analysis

For metabolomics data (n=3 in each experiment) p-value was calculated with a two-tailed Student’s t-test. To evaluate combined significance from independent experiments, p-values were combined with Fisher’s method and corrected for false discovery rate using the Benjamini-Hochberg method. For SA-β-gal activity, p-values were calculated with a Student’s t-test. For cell size analysis, p-values were calculated with a Wilcox-Mann-Whitney test because the data were not normally distributed.

### Immunoblot analysis

Cells were lysed in modified RIPA buffer (50 mM Tris–HCl (pH 7.5), 150 mM NaCl, 50 mM β-glycerophosphate, 0.5 mM NP-40, 0.25% sodium deoxycholate, 10 mM sodium pyrophosphate, 30 mM sodium fluoride, 2 mM EDTA, 1 mM activated sodium vanadate, 20 µg/ml aprotinin, 10 µg/ml leupeptin, 1 mM DTT, and 1 mM phenylmethylsulfonyl fluoride). Whole-cell lysates were resolved by SDS–PAGE on 4– 15% gradient gels and blotted onto nitrocellulose membranes (Bio-Rad). Membranes were blocked for 1 h, and then incubated with primary overnight and secondary antibodies for 2 h. Blots were imaged using the Odyssey Infrared Imaging System (Li-Cor). Primary antibodies used for Western blot analysis were: p16 (10883-1-AP, Proteintech), p21 (2947S, Cell Signaling), PAI-1 (NBP1-19773, Novus Biologicals), γ-H2AX (9718S, Cell Signaling), RRM2 (103193, GeneTex, and HPAA056994, Millipore Sigma), luciferase (L0159, Sigma-Aldrich), hTERT (600-401-252S, Rockland), and anti-beta-actin (66009-1-Ig, Proteintech). Recombinant human p16 protein (Novus H00001029-P01) was used as a positive control for p16 Western blotting.

## ACKNOWLEDGMENTS

This work was supported by The Rose Hills Foundation, The USC Provost’s Office, the USC Translational Imaging Center, and the Viterbi School of Engineering.

## CONFLICT OF INTEREST

The authors declare no conflict of interest.

## AUTHOR CONTRIBUTIONS

AD and NG conceived the study; AD performed cell culture and metabolomics experiments. SP and AD cloned lentiviral vectors for luciferase and hTERT expression. AD, JJ and SF performed and supervised immunofluorescence microscopy. AD, JY, and FS performed Western blotting. AD, SL, JM and PW performed and supervised SA-β-gal activity measurements. AD and NG analyzed all data and wrote the manuscript.

## FOOTNOTES

The abbreviations used are: AMPK, 5’ AMP-activated protein kinase; CAD, carbamoyl-phosphate synthetase 2, aspartate transcarbamylase, and dihydroorotase; CRISPRi, CRISPR-dCas9 interference, dNTP, deoxyribonucleotide triphosphate; EdU, 5-ethynyl-2’-deoxyuridine; FDR, false discovery rate; GSEA, gene set expression analysis; HMEC, human mammary epithelial cells; MSEA, metabolite site enrichment analysis; PD, population doubling; RRM2, ribonucleotide reductase regulatory subunit; SA-β-gal, senescence-associated β-galactosidase; SASP, senescence-associated secretory phenotype; hTERT, human telomerase.

## Notes

#### Summary of Updates

Added further characterization of p21 and PAI-1 upregulation following pharmacological and genetic inhibition of nucleotide synthesis in Fig. 5 and Supp. Fig. 9.

