## Supporting Figures for "Inhibition of nucleotide synthesis promotes replicative senescence of human mammary epithelial cells"

Running title: *Nucleotide synthesis promotes epithelial cell senescence*

To whom correspondence should be addressed: Corresponding author: Nicholas A. Graham, University of Southern California, Los Angeles, 3710 McClintock Ave., RTH 509, Los Angeles, CA 90089. Phone: 213-240-0449;

E-mail addresses: (A. Delfarah), (S. Parrish), (J.A. Junge), (J. Yang), (F. Seo), (S. Li), (J. Mac), (P. Wang), (S.E. Fraser), (N.A. Graham)

**Supporting Fig. 1: Senescent HMEC are frequently multi-nucleated.**

Videos from 3D reconstructions of Z-stack confocal imaging for HMEC cells stained with Hoechst dye (blue) and plasma membrane dye (yellow) for HMEC at PD 10 (A) and HMEC at PD 37 (B, C).

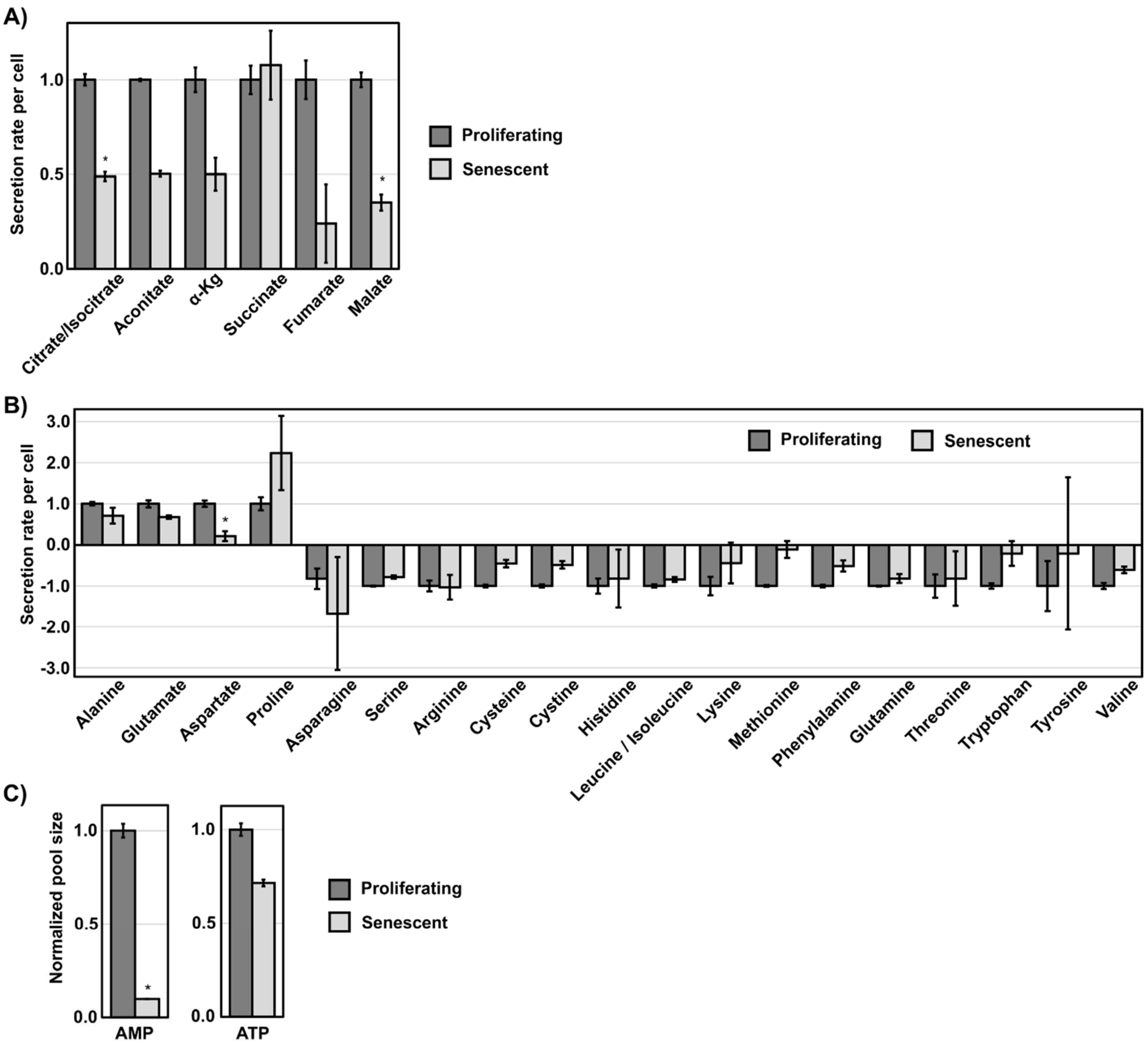


D)

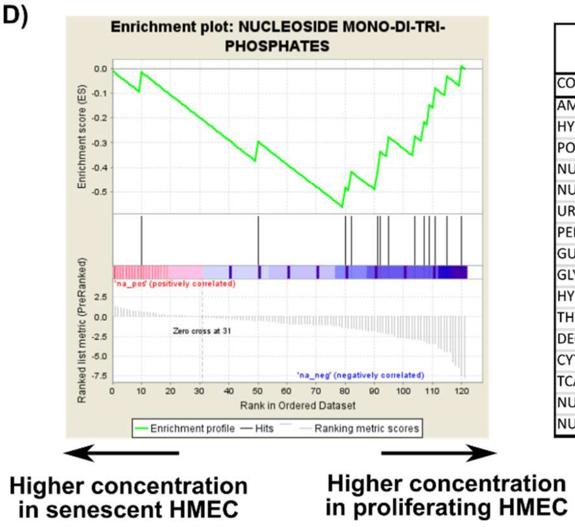


| Replicative senescence pool sizes | | | | |
|---|---|---|---|---|
| COMPOUND CLASS | SIZE | NES | NOM p-val | FDR q-val |
| AMINO ACIDS | 16 | 1.89 | 0.01 | 0.05 |
| HYDROPHOBIC AMINO ACIDS | 5 | 1.87 | 0.01 | 0.03 |
| POLAR AMINO ACIDS | 8 | 1.49 | 0.07 | 0.15 |
| NUCLEOBASES | 6 | 1.36 | 0.14 | 0.20 |
| NUCLEOSIDES | 5 | 1.04 | 0.40 | 0.48 |
| URACIL DERIVATIVES | 5 | 1.03 | 0.40 | 0.41 |
| PENTOSE PHOSPHATE PATHWAY | 6 | -0.69 | 0.85 | 0.86 |
| GUANINE DERIVATIVES | 5 | -0.75 | 0.77 | 0.86 |
| GLYCOLYSIS | 6 | -0.84 | 0.65 | 0.82 |
| HYPOXANTHINE DERIVATIVES | 8 | -0.94 | 0.51 | 0.75 |
| THYMINE DERIVATIVES | 5 | -1.05 | 0.37 | 0.64 |
| DEOXYNUCLEOSIDE MONO-DI-TRI-PHOSPHATE | 10 | -1.52 | 0.06 | 0.13 |
| CYTOSINE DERIVATIVES | 9 | -1.55 | 0.06 | 0.15 |
| TCA CYCLE | 7 | -1.72 | 0.02 | 0.08 |
| NUCLEOSIDE MONOPHOSPHATES | 5 | -1.88 | 0.01 | 0.05 |
| NUCLEOSIDE MONO-DI-TRI-PHOSPHATES | 13 | -2.35 | 0.0004 | 0.003 |

**Supporting Fig. 2: Extracellular medium analysis of amino acid and TCA cycle metabolites, intracellular AMP/ATP levels, and MSEA analysis of senescent HMEC intracellular metabolite pools**
A) Extracellular metabolite secretion data for TCA cycle metabolites in proliferating and senescent HMEC. Metabolite extracts from blank and conditioned media were analyzed by LC-MS. Secretion or uptake values were normalized to integrated cell number. Secreted metabolites have positive values, and consumed metabolites have negative values. * denotes p-value less than 0.05 by FDR-corrected Student's t-test. See Supporting Table S1 for all measured metabolites.
B) Same as in A for amino acids. * denotes p-value less than 0.05 by FDR-corrected Student's t-test. See Supporting Table S1 for all measured metabolites.
C) Intracellular pool sizes of AMP and ATP of proliferating and senescent HMEC. * denotes p-value less than 0.05 by FDR-corrected Student's t-test. See Supporting Table S2 for all measured metabolites.
D) Metabolite set enrichment analysis (MSEA) analysis for intracellular metabolites pool sizes of proliferating and senescent HMEC. Metabolites were ranked based on $\log_2$ fold change of senescent/proliferating. Shown are the mountain plot of nucleoside mono/di/tri-phosphates (left), and table of all tested metabolic pathways (right).

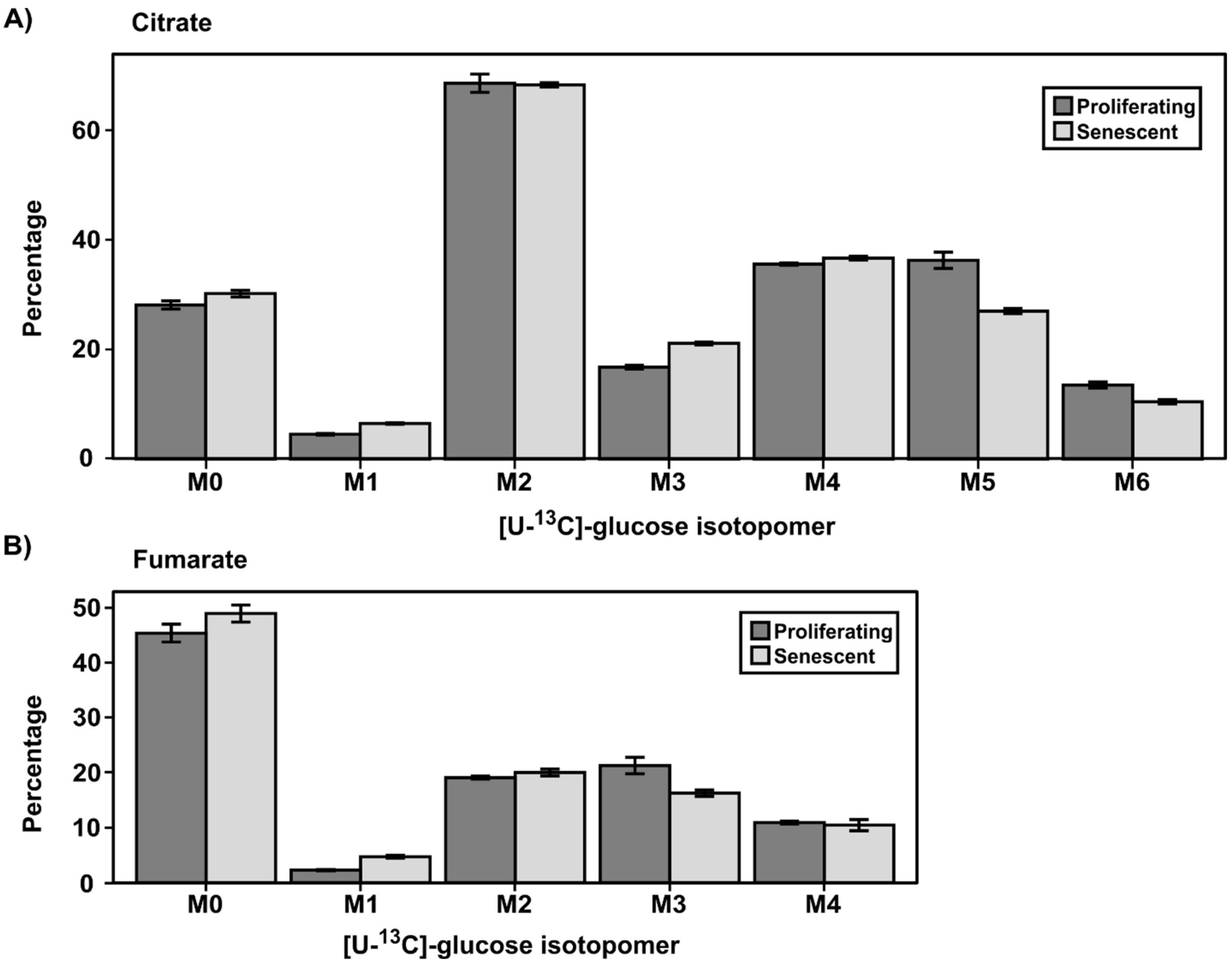


**Supporting Fig. 3: [U-$^{13}$C]-labeled glucose isotopomer distributions are not altered in senescent HMEC.**

A-B) [U-$^{13}$C]-labeled glucose isotopomer distributions of citrate and fumarate. Proliferating and senescent HMEC show similar labeling patterns for TCA cycle metabolites. See Supporting Table S3 for all measured metabolite isotopomer distributions.

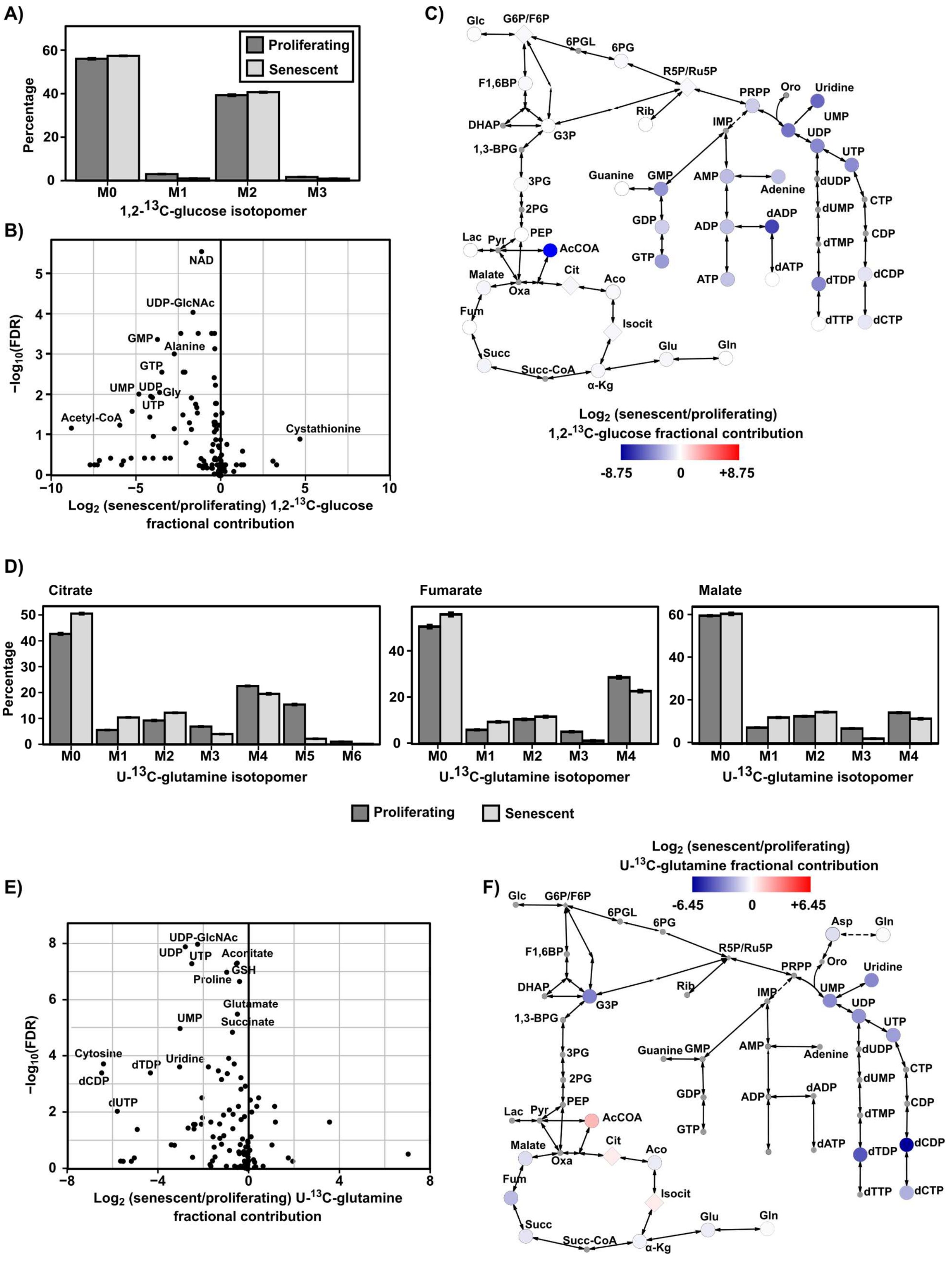

A)
Proliferating
Senescent
Percentage
M0
M1
M2
M3
1,2-$^{13}$C-glucose isotopomer
B)
NAD
UDP-GlcNAc
GMP
Alanine
GTP
UMP
UDP
Gly
UTP
Acetyl-CoA
Cystathionine
−log$_{10}$(FDR)
Log$_2$ (senescent/proliferating) 1,2-$^{13}$C-glucose fractional contribution
C)
Glc
G6P/F6P
6PGL
6PG
R5P/Ru5P
F1,6BP
DHAP
G3P
1,3-BPG
3PG
2PG
PEP
Lac
Pyr
AcCOA
Malate
Oxa
Cit
Aco
Fum
Isocit
Succ
Succ-CoA
α-Kg
Glu
Gln
Rib
PRPP
Oro
Uridine
IMP
UMP
UDP
UTP
Guanine
GMP
AMP
Adenine
GDP
GTP
ADP
dADP
ATP
dATP
dUDP
dUMP
dTMP
dTDP
dTTP
CTP
CDP
dCDP
dCTP
Log$_2$ (senescent/proliferating)
1,2-$^{13}$C-glucose fractional contribution
-8.75
0
+8.75
D)
Citrate
Fumarate
Malate
Percentage
M0
M1
M2
M3
M4
M5
M6
U-$^{13}$C-glutamine isotopomer
Proliferating
Senescent
E)
UDP-GlcNAc
UDP
UTP
Aconitate
GSH
Proline
Glutamate
Succinate
UMP
Cytosine
dCDP
dTDP
Uridine
dUTP
−log$_{10}$(FDR)
Log$_2$ (senescent/proliferating) U-$^{13}$C-glutamine fractional contribution
F)
Log$_2$ (senescent/proliferating)
U-$^{13}$C-glutamine fractional contribution
-6.45
0
+6.45
Asp
Gln
Oro
dCDP
dCTP
dTTP

**Supporting Fig. 4: [1,2-$^{13}$C]-glucose and [U-$^{13}$C]-glutamine stable isotope tracing**

A) [1,2-$^{13}$C]-labeled glucose isotopomer distribution. The ratio of M1 to M2 lactate does not show a significant change between proliferating and senescent HMEC.

B) Fractional contribution of [1,2-$^{13}$C]-glucose to purines and pyrimidines is downregulated in senescent HMEC. Volcano plot represents $\log_2$ fold change (senescent/proliferating) for fractional contribution of [1,2-$^{13}$C]-glucose and FDR-corrected p-value.

C) Metabolic pathway map depicting the average $\log_2$ fold change (senescent/proliferating) of fractional contribution of [1,2-$^{13}$C]-glucose using the indicated color scale. Metabolites that were not measured are shown as small grey colored shapes.

D) [U-$^{13}$C]-labeled glutamine isotopomer distributions for TCA cycle metabolites. The ratio of reductive to oxidative TCA cycle is slightly decreased in senescent HMEC.

E) Fractional contribution of [U-$^{13}$C]-glutamine to pyrimidines is downregulated in senescent HMEC. Volcano plot represents average $\log_2$ fold change (senescent/proliferating) for fractional contribution of [U-$^{13}$C]-labeled glutamine and FDR-corrected combined Fisher's combined p-value from two independent experiments.

F) Metabolic pathway map depicting the average $\log_2$ fold change (senescent/proliferating) of fractional contribution of [U-$^{13}$C]-glutamine using the indicated color scale. Metabolites that were not measured or had less than 3% fractional contribution are shown as small grey colored shapes.

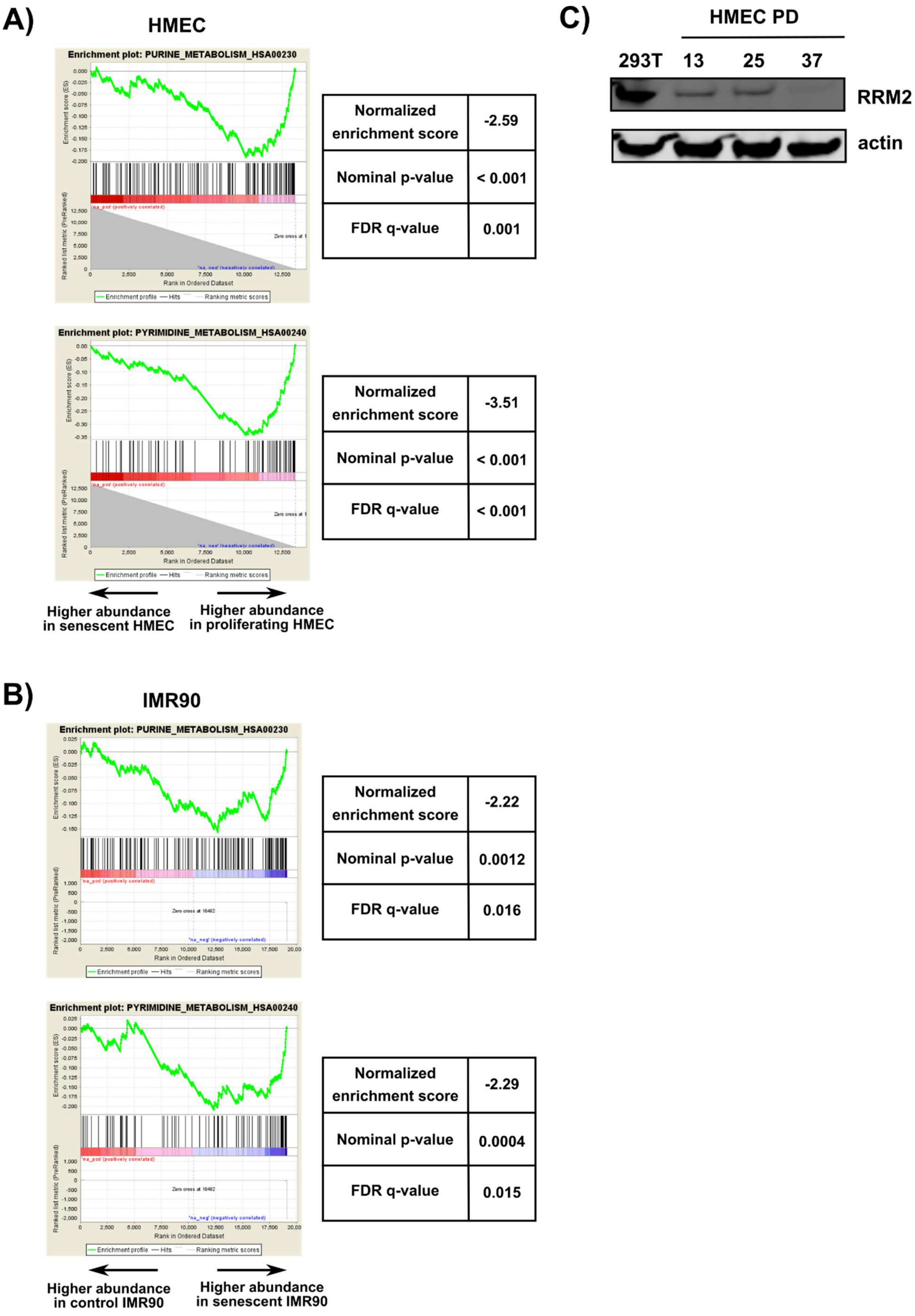
A)
HMEC
Enrichment plot: PURINE_METABOLISM_HSA00230
Normalized enrichment score
-2.59
Nominal p-value
< 0.001
FDR q-value
0.001
Enrichment plot: PYRIMIDINE_METABOLISM_HSA00240
Normalized enrichment score
-3.51
Nominal p-value
< 0.001
FDR q-value
< 0.001
Higher abundance in senescent HMEC
Higher abundance in proliferating HMEC
B)
IMR90
Enrichment plot: PURINE_METABOLISM_HSA00230
Normalized enrichment score
-2.22
Nominal p-value
0.0012
FDR q-value
0.016
Enrichment plot: PYRIMIDINE_METABOLISM_HSA00240
Normalized enrichment score
-2.29
Nominal p-value
0.0004
FDR q-value
0.015
Higher abundance in control IMR90
Higher abundance in senescent IMR90
C)
HMEC PD
293T
13
25
37
RRM2
actin
Enrichment profile
Hits
Ranking metric scores
Rank in Ordered Dataset

**Supporting Fig. 5: RNA expression analysis confirms reduced nucleotide synthesis in senescent cells.** A-B) GSEA analysis of A) microarray data from senescent HMEC (1) and B) RNAseq data from senescent IMR90 cells (2). For HMEC microarray data, genes were compared across pairwise comparisons for stasis and pre-stasis cell cultures. Genes were then ranked by their average rank from individual experiments. For IMR90 RNAseq data, genes were ranked based on signal to noise ratio of senescent/non-senescent. Results show significant suppression of genes in the purine (hsa00230) and pyrimidine (hsa00240) pathways in senescent cells.

C) Western blotting with an RRM2 antibody (Sigma) targeting a distinct epitope from Figure 3E revealed downregulation of RRM2 expression in senescent HMEC. Actin was used as an equal loading control.

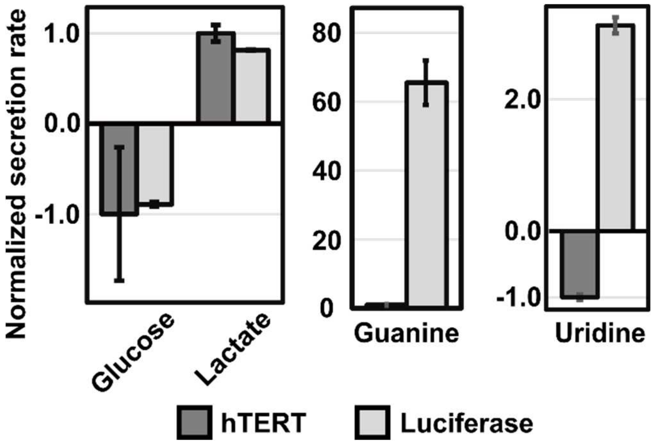


**Supporting Fig. 6 Extracellular metabolites consumption and secretion analysis of luciferase- and hTERT-expressing HMEC.**
Senescent luciferase-expressing HMEC do not exhibit a glycolytic shift but do exhibit increased guanine and uridine secretion. Extracellular metabolite secretion and consumption data was measured by LC-MS using metabolite extracts from blank media and conditioned media from luciferase-expressing and hTERT-expressing cells. Secretion is calculated as (conditioned media – blank media) normalized to integrated cell number. Secreted metabolites have positive values, and consumed metabolites have negative values. At the time of extraction, luciferase-expressing cells but not hTERT-expressing cells were senescent.

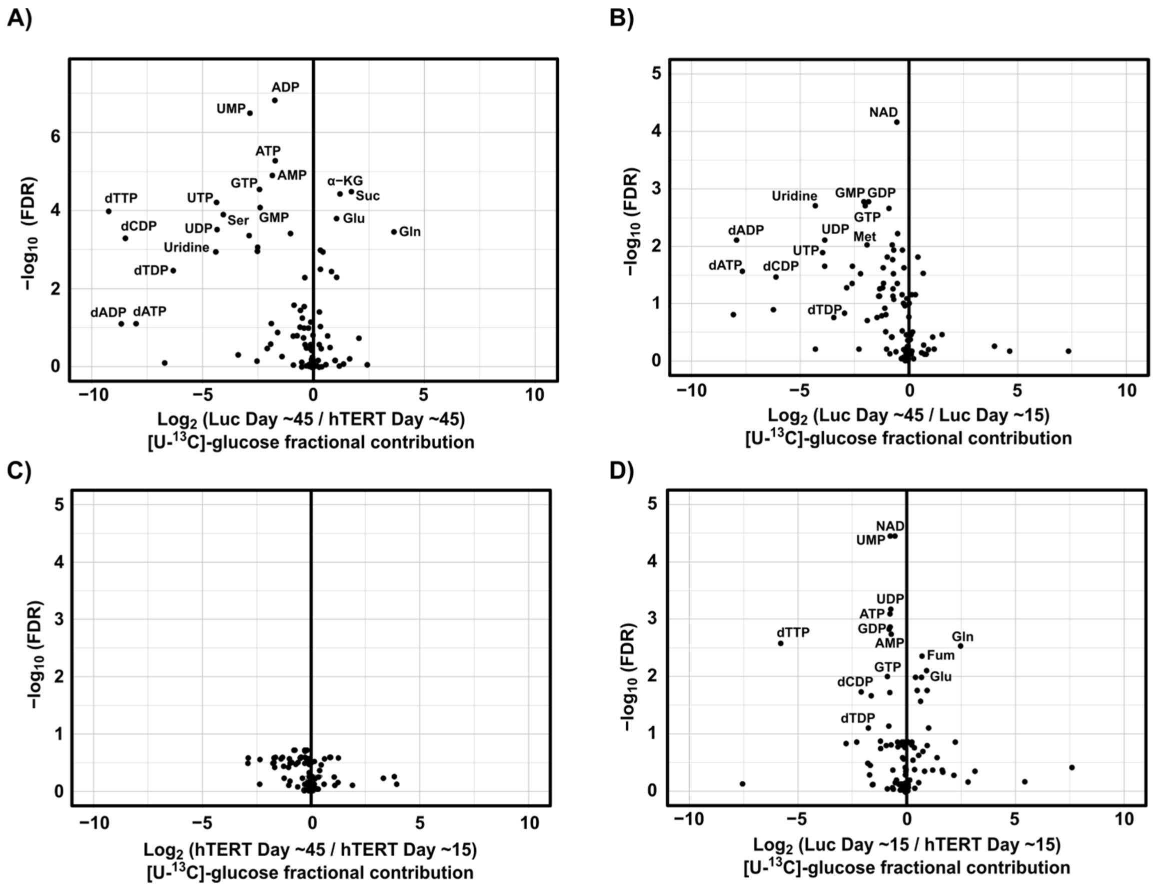


**Supporting Fig. 7: Glucose fractional incorporation analysis of luciferase- and hTERT-expressing HMEC at low and high population doublings.**

Volcano plots representing the average $\log_2$ fold change versus the FDR-corrected p-value of [U-$^{13}$C]-glucose fractional contribution for:

A) Senescent luciferase- (Day ~45) v. proliferating hTERT-expressing HMEC (Day ~45).
B) Senescent luciferase- (Day ~45) v. proliferating luciferase-expressing HMEC (Day ~15).
C) Proliferating hTERT- (Day ~45) v. proliferating hTERT-expressing HMEC (Day ~15).
D) Proliferating luciferase- (Day ~15) v. proliferating hTERT-expressing HMEC (Day ~15).
Days represent the time since completion of drug selection following viral infection.

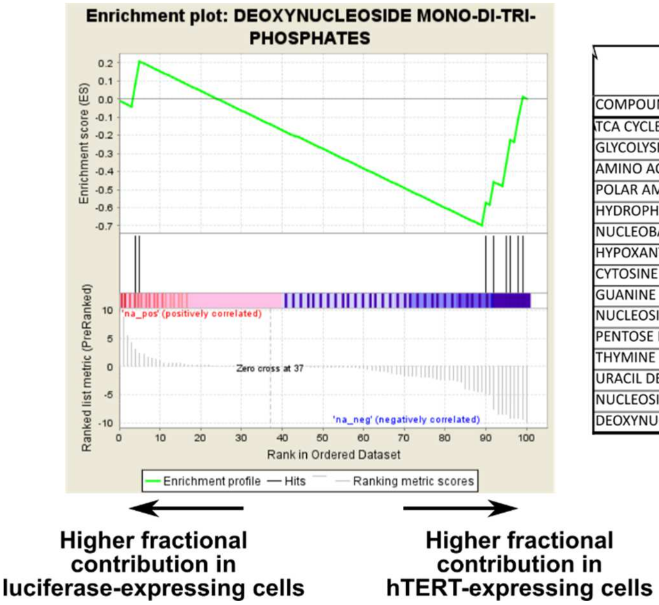


| hTERT immortalization [U-$^{13}$C]-glucose fractional label | | | | |
|---|---|---|---|---|
| COMPOUND CLASS | SIZE | NES | NOM p-val | FDR q-val |
| TCA CYCLE | 7 | 2.45 | 0.001 | 0.001 |
| GLYCOLYSIS | 6 | 1.56 | 0.06 | 0.19 |
| AMINO ACIDS | 16 | 1.22 | 0.22 | 0.52 |
| POLAR AMINO ACIDS | 8 | 1.06 | 0.36 | 0.65 |
| HYDROPHOBIC AMINO ACIDS | 5 | 0.74 | 0.79 | 1.00 |
| NUCLEOBASES | 6 | 0.71 | 0.82 | 0.97 |
| HYPOXANTHINE DERIVATIVES | 8 | 0.65 | 0.89 | 0.89 |
| CYTOSINE DERIVATIVES | 9 | -1.08 | 0.36 | 0.35 |
| GUANINE DERIVATIVES | 5 | -1.10 | 0.31 | 0.37 |
| NUCLEOSIDES | 5 | -1.17 | 0.25 | 0.35 |
| PENTOSE PHOSPHATE PATHWAY | 6 | -1.31 | 0.15 | 0.25 |
| THYMINE DERIVATIVES | 5 | -1.85 | 0.01 | 0.03 |
| URACIL DERIVATIVES | 5 | -2.12 | 0.00 | 0.01 |
| NUCLEOSIDE MONO-DI-TRI-PHOSPHATES | 13 | -2.25 | 0.002 | 0.003 |
| DEOXYNUCLEOSIDE MONO-DI-TRI-PHOSPHATES | 10 | -2.32 | 0.001 | 0.004 |

**Supporting Fig. 8: Metabolite set enrichment analysis of hTERT immortalization.**
MSEA analysis for fractional contribution of [U-$^{13}$C]-glucose in hTERT immortalized HMEC. Metabolites were ranked based on $\log_2$ fold change of luciferase/hTERT.

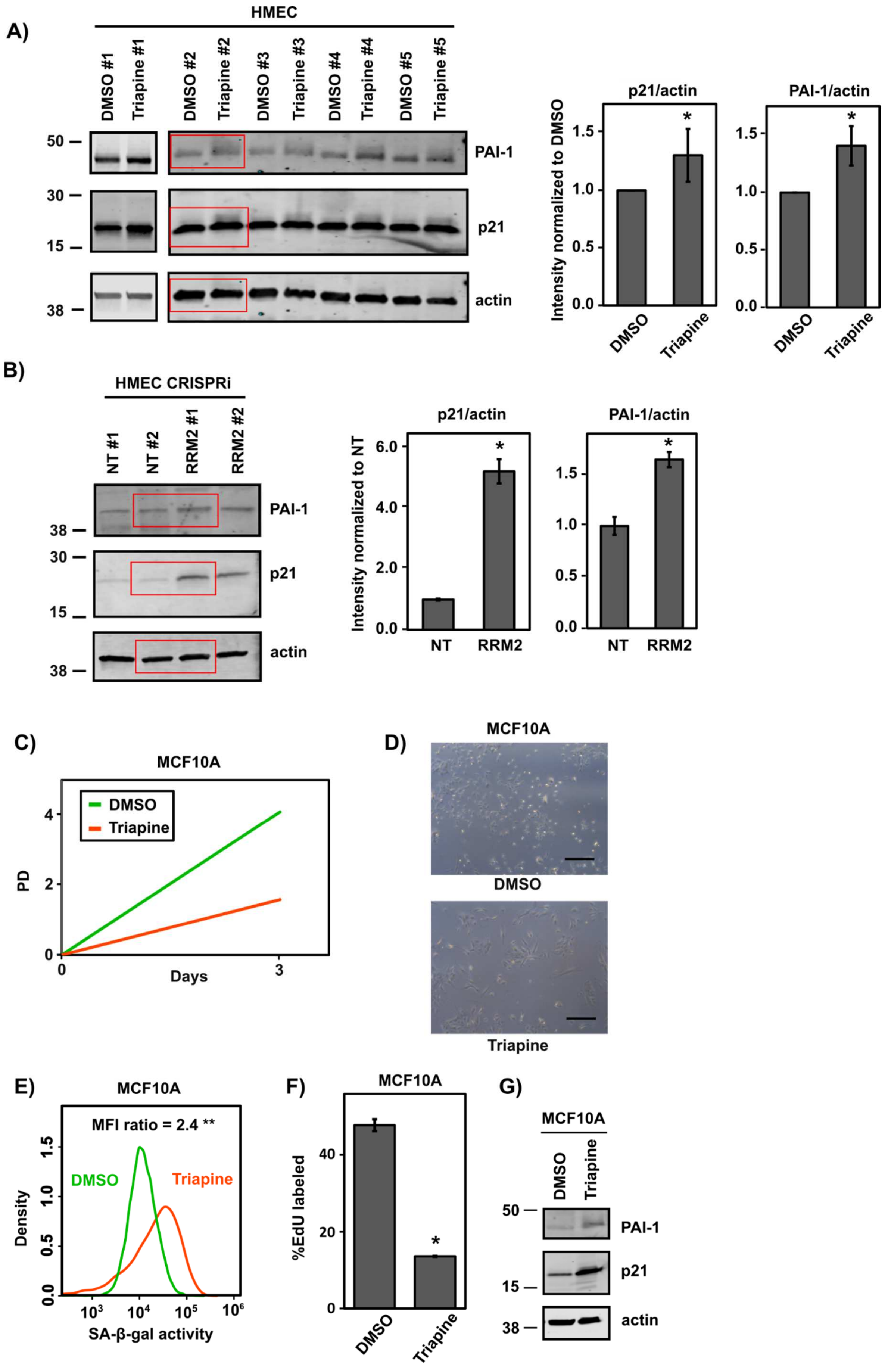
A)
HMEC
DMSO #1
Triapine #1
DMSO #2
Triapine #2
DMSO #3
Triapine #3
DMSO #4
Triapine #4
DMSO #5
Triapine #5
50
30
15
38
PAI-1
p21
actin
p21/actin
PAI-1/actin
Intensity normalized to DMSO
1.5
1.0
0.5
0.0
*
DMSO
Triapine
B)
HMEC CRISPRi
NT #1
NT #2
RRM2 #1
RRM2 #2
38
30
15
PAI-1
p21
actin
Intensity normalized to NT
6.0
4.0
2.0
0.0
NT
RRM2
C)
MCF10A
DMSO
Triapine
PD
Days
0
2
4
3
D)
MCF10A
DMSO
Triapine
E)
MCF10A
MFI ratio = 2.4 **
Density
SA-β-gal activity
$10^3$
$10^4$
$10^5$
$10^6$
F)
MCF10A
%EdU labeled
40
20
0
G)
MCF10A
PAI-1
p21
actin

**Supporting Fig. 9: Inhibition of RRM2 induces senescence in HMEC and MCF10A cells.**
A) Western blots of p21, PAI-1, and actin for triapine-treated HMEC. Blots represent five independent experiments. Red boxes indicate bands that are also shown on Fig. 5E. uantification of bands is represented on bar plots as the average of five independent experiments. * and ** denote p-values of 0.006 and 0.04 by Student's t-test, respectively.
B) Western blot of p21, PAI-1, and actin for HMEC infected with sgRNA against RRM2 or non-targeting control (NT). Blots represent two independent experiments. Red boxes indicate bands that are also shown on Fig. 5I. Quantification of bands are represented on bar plots as average of two independent experiments. * denotes p-value less than 0.045 by Student's t-test.
C) Treatment of MCF10A cells with the RRM2 inhibitor triapine inhibits cell growth. Cells were treated with 2 µM of triapine or DMSO for 72 h.
D) Phase contrast images of MCF10A treated with triapine or DMSO as in A). Scale bar is 100 µm.
E) Triapine-treated MCF10A showed increased activity of SA-β-gal measured by fluorescence signal of $C_{12}$FDG. ** denotes p-value less than 0.0001 by Student's t-test.
F) DNA synthesis measured by EdU incorporation. * denotes p-value less than 0.02 by Student's t-test.
G) Western blot of p21, PAI-1, and actin for DMSO- or triapine-treated MCF10A cells.

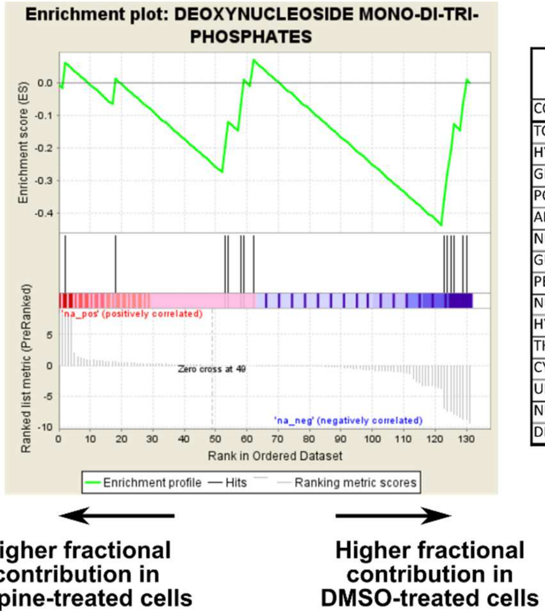


| Triapine-induced senescence [U- $^{13}$C]-glucose fractional label | | | | |
|---|---|---|---|---|
| COMPOUND CLASS | SIZE | NES | NOM p-val | FDR q-val |
| TCA CYCLE | 7 | 1.83 | 0.02 | 0.11 |
| HYPOXANTHINE DERIVATIVES | 8 | 1.15 | 0.28 | 0.97 |
| GLYCOLYSIS | 6 | 1.15 | 0.28 | 0.65 |
| POLAR AMINO ACIDS | 8 | 1.04 | 0.38 | 0.67 |
| AMINO ACIDS | 16 | 1.04 | 0.39 | 0.55 |
| NUCLEOSIDES | 5 | 0.82 | 0.68 | 0.68 |
| GUANINE DERIVATIVES | 5 | -0.78 | 0.73 | 0.74 |
| PENTOSE PHOSPHATE PATHWAY | 6 | -0.85 | 0.64 | 0.72 |
| NUCLEOBASES | 6 | -1.21 | 0.22 | 0.28 |
| HYDROPHOBIC AMINO ACIDS | 5 | -1.22 | 0.21 | 0.31 |
| THYMINE DERIVATIVES | 5 | -1.41 | 0.11 | 0.18 |
| CYTOSINE DERIVATIVES | 9 | -1.74 | 0.02 | 0.06 |
| URACIL DERIVATIVES | 5 | -1.77 | 0.02 | 0.07 |
| NUCLEOSIDE MONO-DI-TRI-PHOSPHATES | 13 | -1.78 | 0.02 | 0.09 |
| DEOXYNUCLEOSIDE MONO-DI-TRI-PHOSPHATES | 10 | -1.82 | 0.02 | 0.15 |

**Supporting Fig. 10: Metabolite set enrichment analysis of triapine-induced senescence.**
MSEA analysis for fractional contribution of [U-$^{13}$C]-glucose in triapine-induced senescence. Metabolites were ranked based on $\log_2$ of triapine/DMSO.
